# Image-based profiling of healthy donor immune cells with Blood Cell Painting reveals novel genotype-phenotype associations

**DOI:** 10.1101/2024.05.17.594648

**Authors:** Caroline Högel-Starck, Veera A. Timonen, Gantugs Atarsaikhan, Isabel Mogollon, Minttu Polso, Antti Hassinen, Jarno Honkanen, Cindy David Sarmento, Jessica Koski, Julius Soini, Lea Urpa, Tanja Ruokoranta, Toveann Ahlnäs, Julianna Juvila, Juho J. Miettinen, Rodosthenis S. Rodosthenous, FinnGen, Outi Kilpivaara, Mikko Arvas, Caroline A. Heckman, Jukka Partanen, Mark Daly, Aarno Palotie, Hanna M. Ollila, Lassi Paavolainen, Vilja Pietiäinen, Esa Pitkänen

## Abstract

The morphological diversity of blood immune cells of healthy individuals, critical for recognizing disease-related phenotypes, remains largely uncharacterized. To address this gap, we developed Blood Cell Painting (BCP): a high content, high throughput fluorescence imaging assay for peripheral blood mononuclear cells. We generated a BCP Atlas with images of 50 million cells from 390 healthy blood donors, identifying 18 distinct immune cell morphology clusters. A genome-wide association study of BCP-derived imaging-based cellular features revealed 93 significant associations across 30 genetic loci. These loci include genes linked to mast cell function, inflammation, immune signaling, mitochondrial maintenance and circadian immune modulation. We also observed correlations between immune cell morphological features and clinical traits, such as respiratory conditions and healthcare visits related to contraceptive management, potentially reflecting hormonal influences on immune cell phenotypes. As a proof of concept for clinical application, acute myeloid leukemia subtypes were distinguished by BCP. Our study establishes BCP as a versatile method for immune cell profiling to uncover genetic, phenotypic and clinical determinants of immune cell morphology in health and disease.

## Background

Despite the success of population-scale genome-wide (GWAS) and phenome-wide association studies (PheWAS) (1), the biological mechanisms linking germline variants to phenotypes remain poorly understood. A crucial intermediate between genotypes and disease phenotypes lies at the level of cellular phenotypes, which can be interrogated at the single-cell level, for example, with single-cell RNA-sequencing (scRNA-seq) and high content imaging (HCI). Single-cell profiling has revealed an unprecedented cellular heterogeneity in healthy and diseased tissues, with implications for the diagnosis and treatment of diseases through an improved understanding of immunity (2). Sequencing-based approaches allow the identification of cell types and subtypes, and mapping of their evolutionary trajectories (3), but these methods are costly and do not capture rich cellular morphology or spatial architecture. By contrast, HCI allows millions of cells to be routinely profiled and quantified in a single experiment, providing cellular context and morphological details at a lower cost (4–6). Consequently, HCI has been used to reveal how immune cell phenotypes reflect molecular and personal health information and how leukemic cancer cells respond, at single-cell level, to different drugs in clinical trials (7,8).

Cell Painting is a widely adopted HCI assay for morphological and functional profiling of single cells and cell organelles via fluorescence microscopy (4,5). Cellular phenotypes derived with Cell Painting have been associated with somatic variants in cancer, transcriptomics, as well as genetic and chemical perturbations (9–11). For example, recent large-scale studies have combined Cell Painting with transcriptomics for >28,000 perturbations (12) and performed >20,000 CRISPR perturbation experiments in >30 million cells, resulting in an atlas of biological pathways and interactions (13). To date, Cell Painting has primarily been applied to study the adherent cancer cell lines and human fibroblasts (4,14), with less focus on human immune cells playing a central role in health and disease.

In this study, we developed the Blood Cell Painting (BCP) workflow (**Fig. 1**) for high content fluorescence microscopy imaging of peripheral blood mononuclear cells (PBMCs), modified from the original Cell Painting assay (4,5). Using BCP, we profiled PBMC samples from 390 healthy blood donors, yielding 50 million imaged cells. We optimized and benchmarked a popular batch effect correction method Harmony for large-scale BCP data. Finally, we associated BCP profiles to genotypes and health registry phenotypes, revealing novel links to immune cell morphology and function.

**Figure 1.**
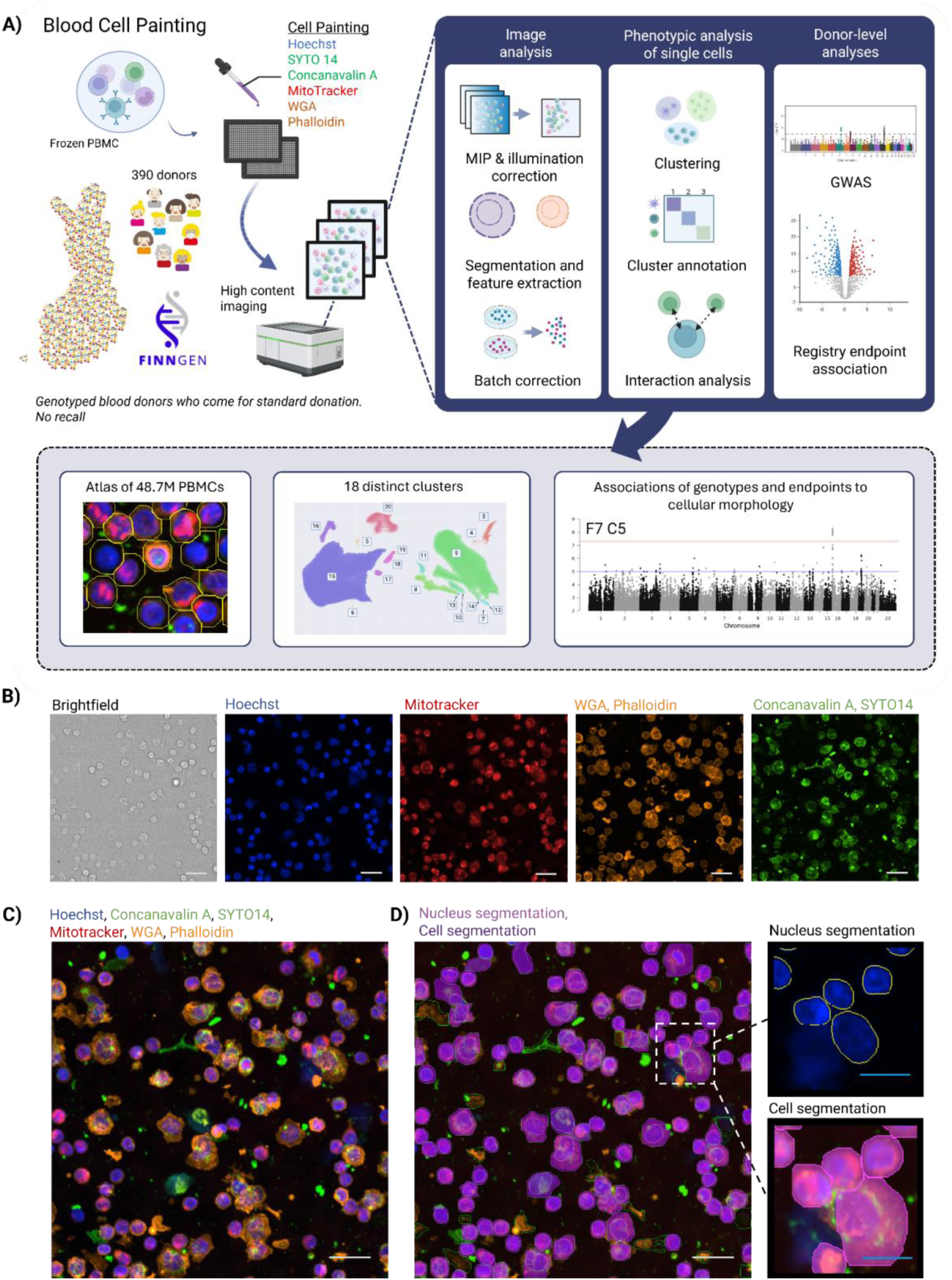
Blood Cell Painting — study design and the BCP assay. A) Overview of the workflow and study results. Created with Biorender.com. B) Example images of PBMCs stained using the BCP assay. C) Example of merged image of four channels containing stains Hoechst 33342, MitoTracker Deep Red, Wheat-germ agglutinin/Alexa Fluor 555 (WGA), Phalloidin/Alexa Fluor 568, Concanavalin A/Alexa Fluor 488 and SYTO14 green fluorescent nucleic acid. D) Visualization of nucleus and cell segmentation. Scale bars: white 20 μm and blue 10 μm.

## Results

### Morphological profiling and cell typing of 50 million PBMCs from 390 healthy blood donors with Blood Cell Painting

We developed a modified Cell Painting protocol for PBMCs, termed Blood Cell Painting (BCP; **Fig. 1 A**; **Methods**). Using the BCP protocol, we profiled PBMCs in 400 samples from 390 healthy blood donors. For each sample, we aimed to image at least 100,000 cells distributed across five wells on the same plate. Altogether, 41 plates were imaged, yielding a total of 50 million cells in 4.74 million images after excluding control samples. We acquired confocal images with four fluorescent channels and widefield images with five fluorescent channels using six fluorescent dyes (**Fig. 1 A**; **Methods**) to label different components of the cell. In addition, brightfield images were collected in both imaging modalities (**Fig. 1 B-D**). A total of 186 features were extracted from images for downstream analyses.

To identify major PBMC cell types in our BCP dataset, we clustered single-cell morphologies from 20 donors and performed supervised cell type annotation where two experts labeled a subset of the cells to train a neural network classifier (**Methods**). Cells were assigned into four classes: 1) monocytes, 2) macrophages, 3) B and T cells, and 4) uncategorised (**Fig. 2 A**). We then evaluated our annotated training set with ten-fold cross-validation (**Methods**). This procedure resulted in cells classified as B and T cells (38% of all cells), monocytes (16%), macrophages (14%) and uncategorised cells or debris (32%) (**Fig. 2 B, C**; **Suppl. Fig. 3 B**; **Suppl. Fig. 2**).

**Figure 2.**
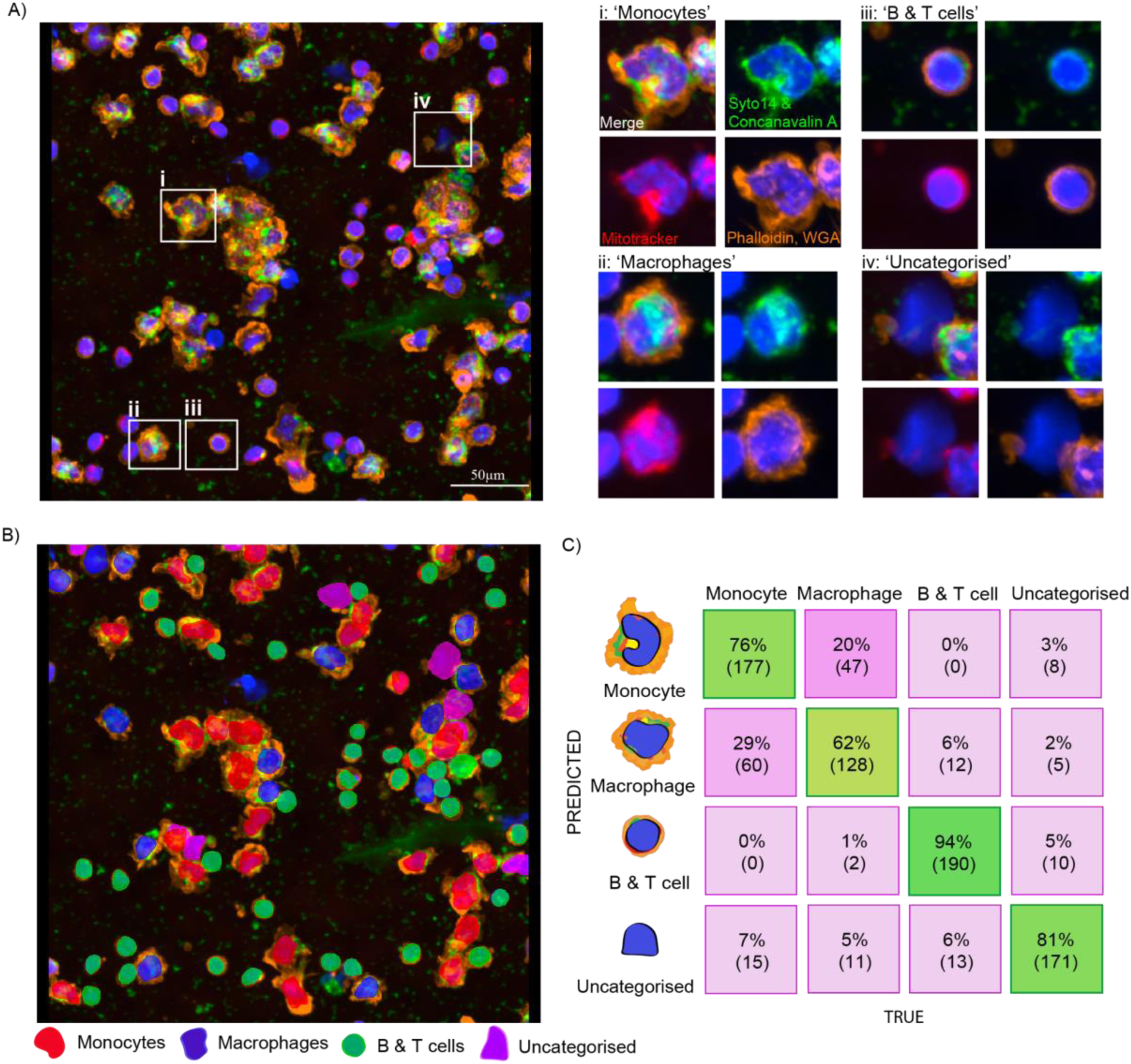
Cell classes based on morphology by expert annotations. A) Examples of cell classes annotated for the training set. Cells were manually annotated into four classes: monocytes (i), macrophages (ii), B & T cells (iii), and uncategorised (iv). B) Single-cell predictions from a representative field-of-view. Classification of cells into four classes is highlighted with the predicted class color: monocytes (red), macrophages (blue), B & T cells (green), uncategorised (magenta). C) Cross-validation confusion matrix obtained for the cell type classifier trained in BIAS software after 10-fold cross-validation. Correct predictions are shown in the diagonal and in green color. Scale bars: 50 µm.

To obtain independent estimates of cell type frequencies, we performed flow cytometry-based immunophenotyping on the same 20 samples (**Methods**; **Suppl. Table 5**). In contrast to the BCP-based typing, flow cytometry indicated B and T cells to be two-fold as frequent (87%; 76% CD3+, 11% CD19+), and five-fold fewer monocytes (3%) (**Suppl. Fig. 3 B, D**; **Suppl. Fig. 4**). Among T cells (CD3+), 23% were CD8+ cytotoxic T cells. NK and dendritic cells were rare (1% and 0.004%, respectively). Despite these differences, both BCP and flow cytometry data showed clear clustering in UMAP embeddings (**Suppl. Fig. 3 A, C**). Notably, many of the cell types that remained uncategorized in BCP formed distinct clusters, suggesting they correspond to specific cell types, dead cells or debris (**Suppl. Fig. 5**). In general, the cell type fractions observed in flow cytometry corresponded to proportions reported in healthy individuals with the same methodology (15). Cell size estimates derived from BCP and flow cytometry (**Suppl. Fig. 3E–F**) reflected known size differences among PBMC subtypes (16,17). These observations across platforms further support the validity of our cell type assignments using BCP. Finally, we found that cell type proportions in BCP were moderately correlated with flow cytometry for both B & T cells and monocyte/macrophage populations (*r*=0.31; **Suppl. Fig. 3G,H**), indicating general concordance between the two methods despite biology- or platform-specific biases.

### Benchmarking batch effect correction methods for Blood Cell Painting profiling data

We next examined the quality and batch effects in the full BCP dataset of 50 million cells. Each donor contributed a median of 119,008 cells, and each well contained a median of 23,682 cells (**Fig. 3 A, B**). Cellular features extracted from the BCP confocal images for the whole dataset of 390 donors revealed a degree of clustering by plate (**Fig. 3 D**), even after normalizing features for controls (**Methods**). This prompted us to quantify batch effects as the degree of unexpected co-occurrence of cells from the same plate (batch effect score; **Methods**). We evaluated the data for batch effects before and after correction using Harmony (18), ComBat (19,20) and linear regression. As batch factors to be corrected in the data, we used plate, well and field-of-view (FOV) separately as well as combinations of these three factors.

**Figure 3.**
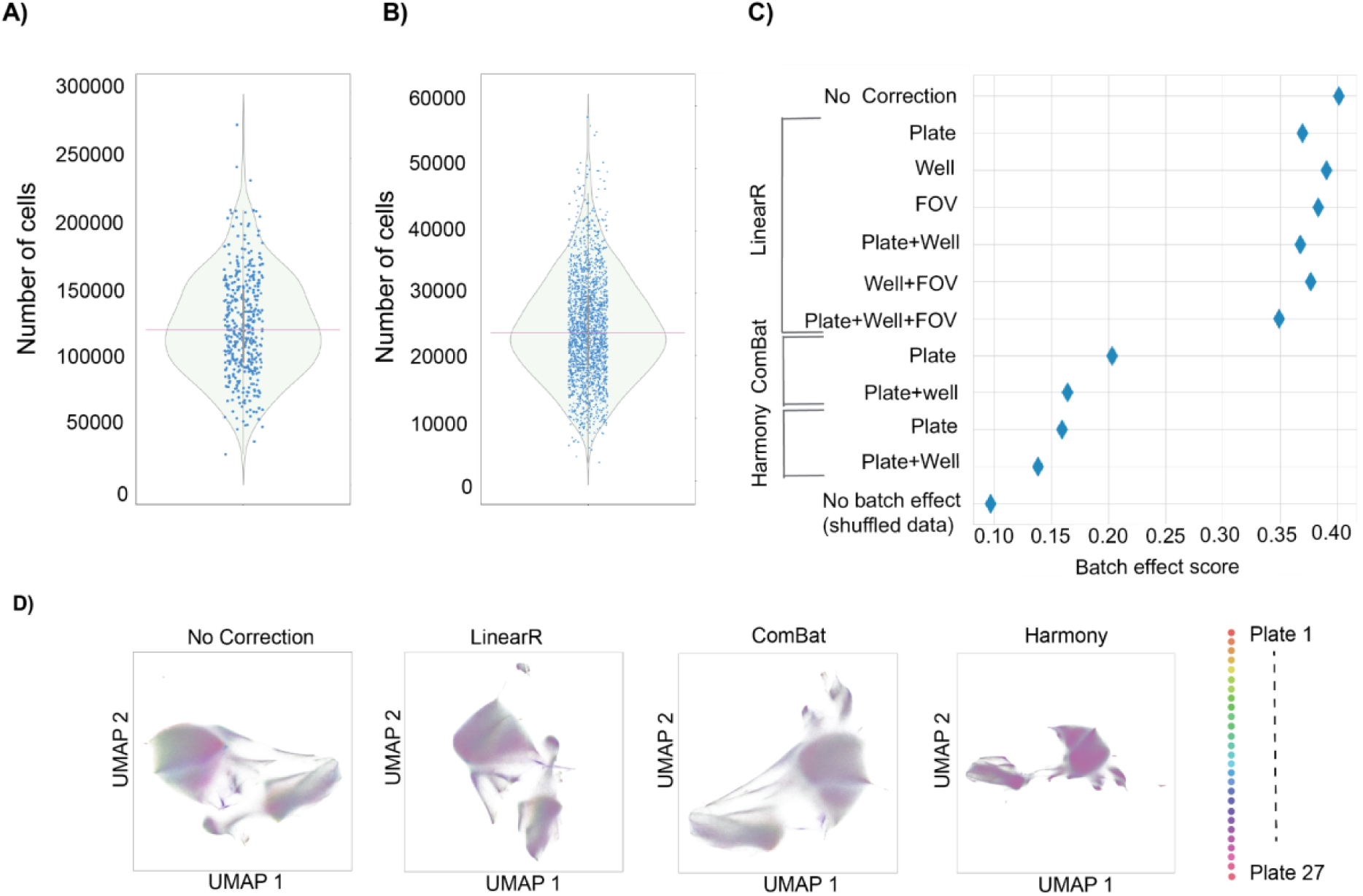
Quality control, and batch effect quantification and correction for BCP data. Cell count distributions of A) 390 donors and B) 2000 wells (4,870,934 cells in total). C) Quantification of batch effect in uncorrected data, and in data corrected with linear regression (LinearR), ComBat, and Harmony for combinations of three batch factors (Plate, Well, FOV). 95% confidence intervals indicated with bars. D) Two-dimensional UMAP representations of cellular features before and after correction with the three methods. Colors indicate the source plate of each cell (27 plates in total).

We evaluated the performance of each method with a subset of the whole dataset containing 2.69 million cells from 269 donors in 27 plates (10,000 cells from each donor). Harmony outperformed the other methods when both plate and well were used as the correcting factors, decreasing the batch effect score by 65.5% compared to 59.1% of ComBat and 8.4% of linear regression (**Fig. 3 C**; **Suppl. Table 6**). Consistently, two-dimensional UMAP representations of cellular features showed minimal clustering by plate in data corrected with Harmony (**Fig. 3 D**). Although we saw improvement when using both plate and well as factors in Harmony and ComBat, we could not evaluate these methods with all three factors (plate+well+FOV) due to excessive memory requirements. Harmony batch-corrected data was used for all subsequent analyses.

### Blood Cell Painting Atlas of peripheral blood mononuclear cells from healthy donors

We combined all 390 donor samples after batch correction to create, to our knowledge, the largest morphological atlas of PBMCs. Cell clustering by morphological profiles yielded 18 distinct clusters encompassing a total of 42.5 million cells (clusters C3-C20; **Fig. 4 A**; **Suppl. Table 8**; **Methods**). Applying our morphology-based cell type classifier to the BCP Atlas revealed the two largest cell clusters to be primarily composed of lymphocyte cells (C15, 58% of all cells), and monocytes and macrophages (C9, 29%), respectively (**Fig. 4 B**). Remaining smaller clusters consisted mostly of uncategorized cells (13%).

**Figure 4.**
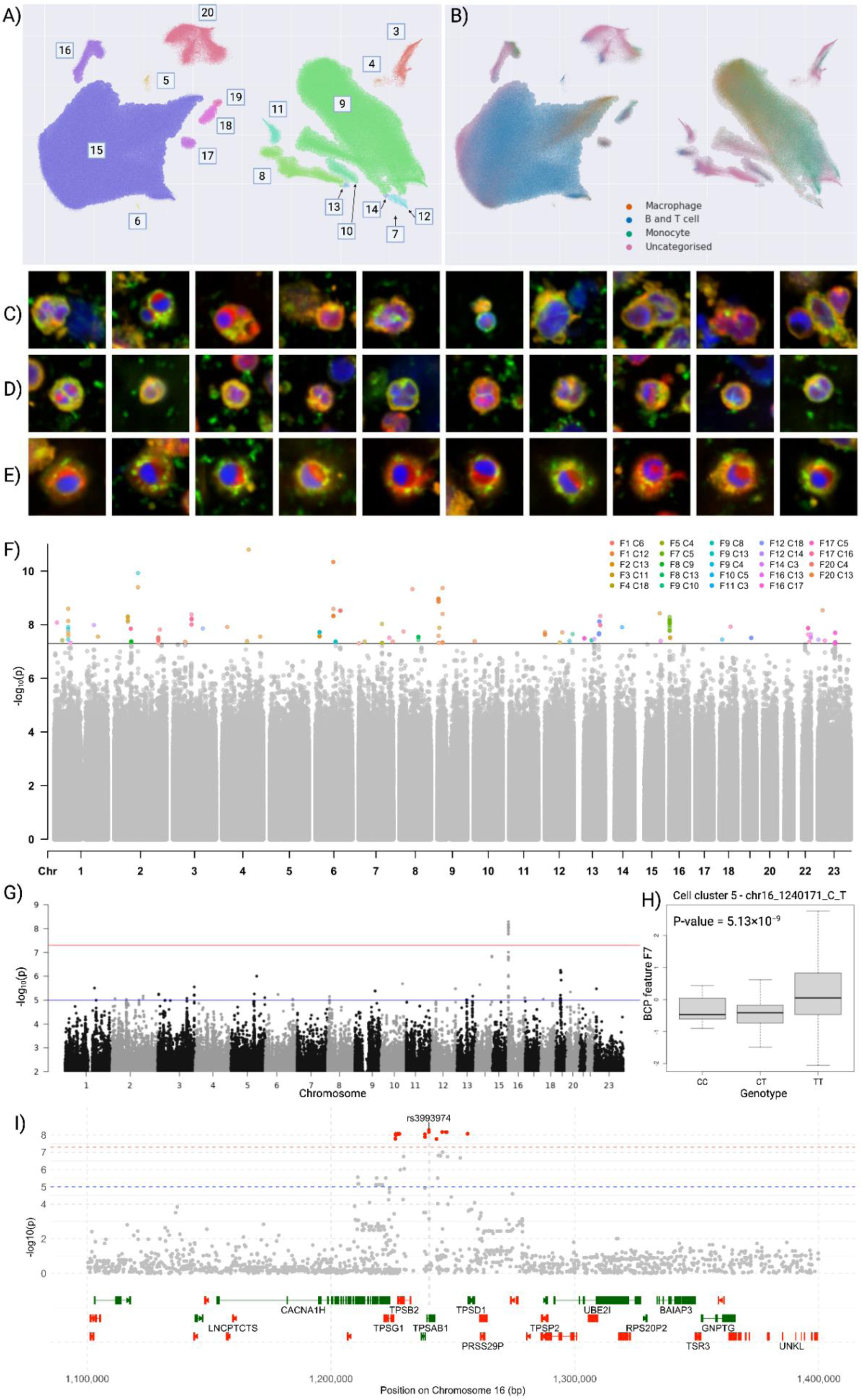
GWAS reveals distinct cell phenotypes are significantly associated with variants. UMAP of the BCP Atlas colored by A) cell clusters C3-C20 and B) cell type identified by a supervised classification procedure. Cell image examples from clusters C) C3, D) C5 and C) C13. F) A combination Manhattan plot of all 93 genome-wide significant associations (-log_10_(p)), where all significant associations for BCP features and clusters are highlighted. G) Genotype-phenotype associations for phenotype F7 in cell cluster C5 (-log_10_(p)), H) Association between lead variant rs3993974 and phenotype F7 in cell cluster C5. I) A locus zoom plot at rs3993974.

Visual examination of BCP images of cells from each cluster confirmed marked morphological differences between clusters (**Fig. 4 C**, **Suppl. Fig. 5**). Several clusters were enriched for cells with C-shaped or otherwise non-spherical nuclei (C3–6, C9, C18, C19). Cluster C11 consisted of cells with elongated shapes, potentially due to interacting cells or active cell division. Intriguingly, cells in cluster C13 showed a striking semicircular pattern on the mitochondrial channel wrapping around a large portion of the nucleus. This pattern may reflect perinuclear mitochondrial aggregation, a localized accumulation of mitochondria around the nucleus (21).

To further delineate the composition of BCP clusters, we correlated their donor frequencies with cell type counts obtained from snRNA-seq for 315 donors (22) (**Suppl. Fig. 6A, Suppl. Table 9**) and with complete blood counts (CBC) for 321 donors (**Suppl. Fig. 6B, Suppl. Table 9; Methods**). The strongest correlations were observed between clusters C9 and monocytes (CD14+, *r*=0.43; CD16+, *r*=0.22), and C15 and lymphoid cells (CD8 naive T, *r*=0.25; B naive, *r*=0.23; CD4 naive T, *r*=0.27), consistent with their morphology and supervised classification (**Fig. 4B, Suppl. Fig. 3G,H**). Based on cell morphology and snRNA-seq data, we hypothesize that C18 may contain non-polarized naive T cells with deep nuclear envelope invaginations (NEIs, “stripy” phenotype; **Suppl. Fig. 5 C18**; CD8 naive *r*=0.13) as recently described (23). Cluster C17 correlated with lymphocytes (B-lymf *r*=0.22, L-lymf *r*=0.24), and memory B cells (*r*=0.15), CD8 naive (*r*=0.14) and CD8 proliferating cells (*r*=0.14), suggesting that this cluster may consist of B and T cells. In cluster C12, we observed larger, possibly damaged or dying cells with strong actin staining (**Suppl. Fig. 5 C12**), with highest correlation to snRNA-seq and CBC monocytes.

We next examined whether cell-cell interactions could be observed in BCP. For each cell, we quantified the relative enrichment (RE) of neighboring cells of each type within a 22 µm radius (**Methods**). We found that each cell type was significantly enriched in neighbors of the same type (RE>1; *p*<1x10^-15^, one-way ANOVA; **Suppl. Fig. 7**). In contrast, other cell types were less frequent within the interaction radius (RE<1). We also quantified the RE in each of the 18 cell clusters and observed that monocytes had a RE slightly above 1 for cluster C9 cells (**Suppl. Fig. 5**), which were earlier classified mostly as monocytes and macrophages (**Fig. 4**). Similarly suggestive of self-enrichment, B/T cell interactions were found to be slightly more frequent in cluster C15 (**Suppl. Fig. 5**).

### Genome-wide association study of Blood Cell Painting features reveals 30 loci influencing immune cell morphology

To discover germline genetic variants influencing immune cell morphology, we performed a GWAS on BCP image-derived features in healthy blood donors. As quantitative traits, we used twenty principal components of donor-averaged BCP features (F1-F20; **Methods**) in 17 cell clusters, capturing major axes of morphological variation, to discover associations of cellular characteristics with germline genetic variants. We analyzed ∼10 million variants (BCP donor minor allele frequency, MAF>1.25%, at least ten carriers in donors) in 368 genotyped donors (FinnGen R12) (1). This yielded 106 genome-wide significant associations (*p*<5x10^-8^) (**Suppl. Fig. 8,9**), of which 93 passed quality control (**Methods**), spanning 30 genomic loci (**Fig. 4 f**, **Suppl. Table 10).** Fine-mapping with SuSiE (24) implicated high-confidence variants in genes related to transcriptional regulation (*C1orf21, ZMYM2*), inflammatory signaling and tissue remodeling (*CHI3L1*), cell cycle and circadian regulation (*CSNK1E*), vesicle trafficking and secretion (*PTPRN2, NCALD*), and extracellular matrix interactions (*PDGFRL*) (**Suppl. Table 13; Methods**).

A prominent association was found in cluster C5 at a locus containing tryptase-encoding genes *TPSB2*, *TPSAB1* and *TPSD1*. The lead variant (rs3993974, GWAS *p*=5.13x10^-9^; SuSiE PIP=0.11; **Suppl. Table 11**, **Suppl. Table 13, Fig. 4 h, i**) is strongly associated with increased tryptase levels in plasma (tryptase µg/l, *β*=0.23, *p*=4.63x10^-11^; **Methods**; **Suppl. Table 12**). This region had a strong expression and splicing quantitative trait locus (eQTL, sQTL) signal for tryptase genes (**Suppl. Table 14**). In addition, this region was marked by direct transcription factor binding by CTCF and nine additional transcription factors, as assessed by ChIPseq (**Suppl. Table 15**). Tryptases are involved in asthma and other allergic and inflammatory disorders, and are contained in mast cells (25). The associated BCP feature (F7) reflected cell length and nuclear stain intensity (**Fig. 4 g, Suppl. Fig. 9 a, e, i; Suppl. Fig. 10).** Cluster C5 also showed associations for nuclear morphology (F17) with rs71330739 (*p*=4.32×10⁻⁸) near cell adhesion and migration related *PARVG* and *PARVB*, and rs62486739 (*p*=3.07×10⁻⁸; PIP=0.44) in *SSBP1*, a housekeeping gene involved in mitochondrial biogenesis (**Suppl. Fig. 9 b, f, j; Suppl. Fig. 10)**. These variants further support the notion that C5 includes rare or immature myeloid populations with distinct nuclear morphologies. While mature mast cells are not generally present in peripheral blood, we speculate that this cluster may be composed of mast or granulocyte cell precursors, or neutrophils (CBC B-neut *r*=0.2; **Suppl. Fig. 6**).

In cell cluster C3, nucleus solidity, cavity and textural patterns (F14) were associated with rs965618 (*p*=2.28x10^-8^; PIP=0.35) (**Suppl. Fig. 9 c, g, k; Suppl. Fig. 10)**. The lead variant is an expression quantitative trait locus (eQTL) across several tissues for *SCO2*, *CPT1B* and *CHKB* that are central regulators of immunity and lipid metabolism across multiple tissues (**Suppl. Table 14**). The locus was also strongly associated with increased mean corpuscular volume (Kanta *β*=0.02, *p*=4.0x10^-27^) and monocyte counts (OpenTargets *β*=0.01, *p*=3.99x10^-8^; **Suppl. Table 12**). In cell cluster C13, which harbored a striking phenotype possibly due to perinuclear mitochondrial aggregation discussed above (**Fig. 4e),** nuclear, mitochondrial and actin characteristics (F8) were associated with rs13251546 with a clear additive effect (**Suppl. Fig. 9 d, h, l; Suppl. Fig. 10)**. The region has previously been associated with visual disorders, varicose veins and smoking status, as well as vision issues (26).

As an example of variants previously implicated in immune-related conditions and highlighted by the BCP GWAS, we found psoriasis-linked variants (OpenTargets *β*=-3.6, *p*=5.00x10⁻¹⁵; **Suppl. Table 12**) at an enhancer site near *CD28* to associate with BCP features (F20; rs115029131, *p*=3.21x10⁻⁸). Many cells in the associating cluster C4 had a non-spherical nucleus, and might thus be monocytes or macrophages (**Suppl. Fig. 5**). Normally CD28 is not expressed in monocytes and macrophages, but can be induced under chronic inflammation (27).

Finally, we performed GWAS for all BCP phenotypes without cell clustering, but no genome-wide significant associations were found. A comparison of our results to previous GWASs (28–30) revealed positive correlations in association strengths (*r*=0.06–0.18, *p*<0.005) (**Suppl. Fig. 11**, **Suppl. Table 16, Suppl. Table 20**).

### Association of BCP cell characteristics to donor age, sex and health registry endpoints

We examined how demographic factors and health registry phenotypes relate to BCP morphological characteristics. Donor age and sex were found to associate (FDR<5%; **Methods**) with several features such as nucleus, mitochondria and RNA intensity and texture as well as cell shape (F1, F2, F7, F13, F16, F17, F20; **Suppl. Table 18**) in cell clusters C9 and C11 (**Suppl. Table 17**). C9 is composed mostly of monocytes and macrophages with strong actin intensity (**Suppl. Fig. 5**). In C11 consisting of elongated cells, sex was associated with cell shape, texture and length as well as RNA dye intensity (FDR=3%; F2).

Morphological features of cells in C19 were associated (FDR<10%; F5, F9, F17) with inflammatory and allergic endpoints such as asthma, sinusitis and pneumonia (**Suppl. Table 19**). These cells were also associated with abnormalities of breathing, septum deviation and vocal larynx issues. Nuclei in C19 exhibit a lobular structure, suggesting they may be granulocytes or their precursors that survived PBMC isolation. Morphological characteristics of cells in C8, potentially composed of T cells, were linked to common rhinitis (F13). Interestingly, healthcare visits related to contraceptive management (ICD-10 Z30) were associated with elongated cell shapes (F16) in C11 (**Suppl. Fig. 5**).

### BCP profiling of acute myeloid leukemia PBMC patient samples

As a proof-of-concept study to test the generalization of the pipeline developed for healthy PBMCs, we performed the BCP protocol on PBMC samples of six acute myeloid leukemia (AML) patients and an AML cell line (MOLM-13) (**Methods**, **Suppl. Table 7**). PBMCs from AML patients displayed distinct morphological profiles (**Fig. 5**). In AMLs with minimal maturation (FAB M1, AML 2-3) and acute monocytic leukemias (M5, AML 6), we observed a cell cluster not found in healthy cells nor the other AML samples. AML samples 4 (M1) and 5 (M5) had low levels of clinically malignant cells in the blood, indicated by most of their cells clustering among the healthy cells. Finally, the AML cell line (MOLM-13) forms distinct clusters within a supercluster composed of monocytes.

**Figure 5.**
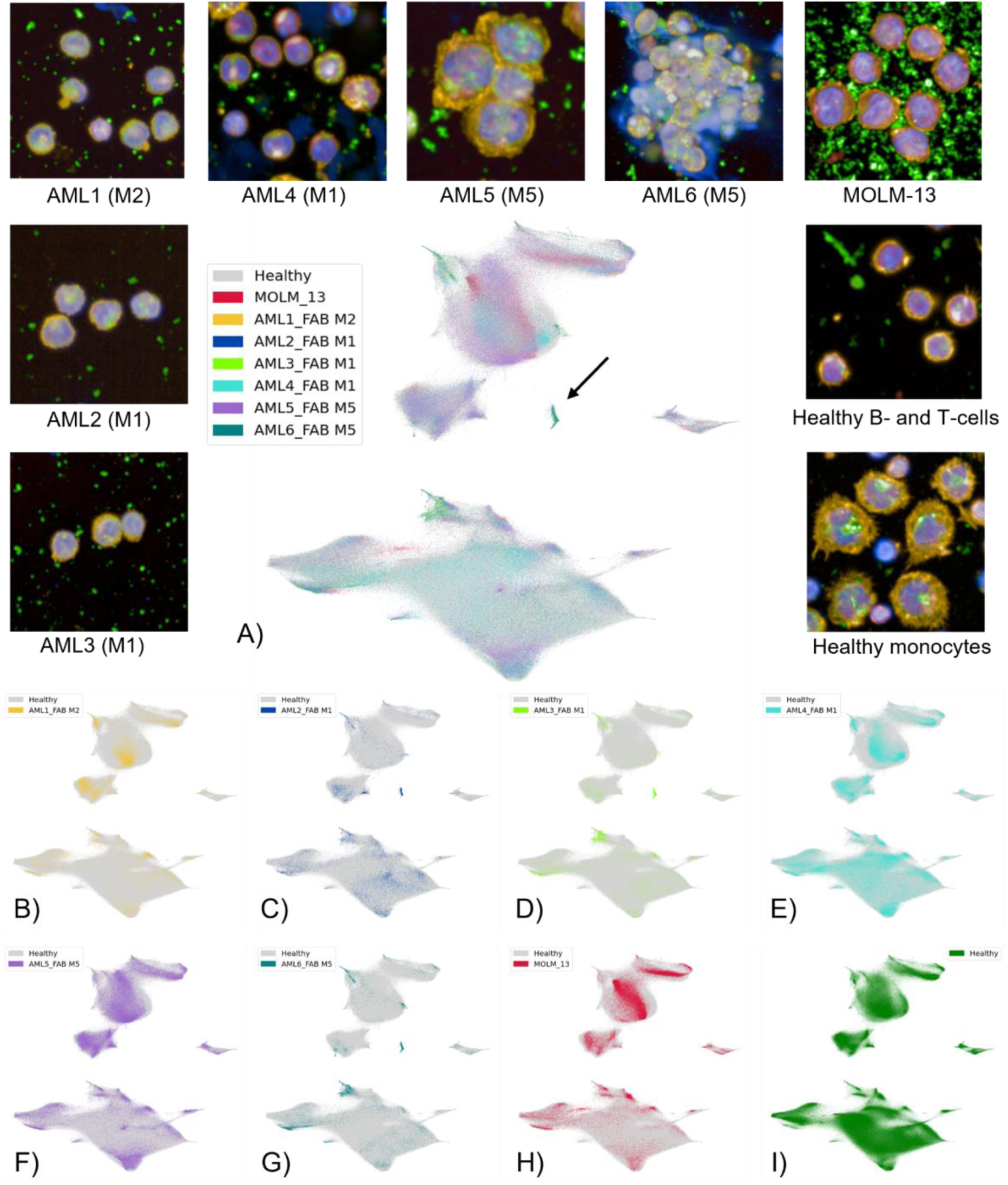
AML samples show a distinct distribution of BCP features compared to healthy donors. A) UMAP of BCP cellular features from AML patient and cell line samples with health PBMCs. Example images of each patient and cell line samples, as well as images of healthy B and T cells and monocytes. A cell cluster not found in healthy cells is indicated with an arrow. B-G) AML patient samples 1-6 and (H) the MOLM-13 sample compared to (I) healthy PBMCs.

## Discussion

In this study, we aimed to characterize the morphological variability of mononuclear cells in healthy blood donors. To achieve this, we developed a modified Cell Painting (4) protocol termed Blood Cell Painting (BCP), specifically for PBMCs, optimizing it for robustness and throughput. With BCP, we created the largest Cell Painting dataset of PBMCs to date, a BCP Atlas encompassing 50 million imaged cells from 390 healthy individuals. The large number of cells profiled allowed us to capture rare morphological phenotypes, such as perinuclear aggregation of mitochondria (31) in immune cells. This work adds a novel dimension to the study of the human immune system through blood profiling (32).

HCI has been shown to exhibit technical variation stemming from experimental conditions such as plate effects and cell handling (33,34). In line with earlier studies (33,35), batch correction with Harmony outperformed alternatives, and we adapted it to scale across tens of millions of cells. Additionally, we observed biases in cell type representation, where monocytes and macrophages appeared more common than expected. The observed difference in cell frequency between flow cytometry and the BCP protocol likely stems from two related factors: higher concentration of B/T cells at the edges of the well and the non-adherent nature of these cell types (36), making them more susceptible to loss during washing compared to monocytes. Further optimization of Poly-L-lysine coating of the wells to improve cell adhesion can mitigate this issue. Our supervised cell type assignment was able to differentiate between lymphoid and myeloid populations. Marker-based sorting could improve cell typing accuracy for rare populations such as NK and dendritic cells.

Correlating donor-level BCP cluster frequencies with cell type counts from snRNA-seq and complete blood counts provided independent support for our morphology-based cell type classification. Concordant patterns observed for myeloid and lymphoid populations indicate that variation captured by BCP reflects true biological differences in circulating immune cell composition rather than technical artefacts. These cross-platform correlations further demonstrate BCP’s capacity to capture physiologically meaningful immune cell heterogeneity.

Driven by the genotyping and health registry data available in FinnGen (1) for our study cohort, we were able to examine the determinants and correlates of immune cell morphological phenotypes. We discovered 30 novel genomic loci associated with morphological cell characteristics. The genetic architecture of immune cell morphology reflected variation in immune cell activation and differentiation, subcellular structural organization, and susceptibility to allergic and inflammatory conditions. As an example, we uncovered genetic variation at tryptase-encoding genes to strongly influence morphology of a distinct cluster of cells. Tryptases, the most abundant serine proteases stored in mast cell granules, are vital for mast cell function and activation, with a key role in allergic reactions and asthma (25). While mature mast cells are not typically found in peripheral blood, immature mast cells can be present.

We also observed associations between BCP features and genetic variants previously implicated in psoriasis. For instance, a psoriasis-linked variant near *CD28* was associated with cells characterized by non-spherical nuclei. This morphology may reflect, for example, activated or differentiating monocytes and macrophages. These cells are known to undergo nuclear deformation in response to environmental stimuli and immune activation (39). Systemic inflammation in psoriasis has been associated with elevated levels of low-density granulocytes (LDGs), a neutrophil subset with segmented or irregular nuclei that co-fractionate with PBMCs (40). These findings suggest that inflammatory skin disorders may leave a morphological footprint in the peripheral immune system, linking genotype-associated immune dysregulation with altered nuclear architecture.

Previous studies have identified numerous germline variants associated with immune cell traits measured by flow cytometry (28,29,41–44) and imaging (30), and immunologic endpoints (45). Although the genome-wide significant variants found in this study were novel, we found variants with *p*<10^-5^ in our study that had been previously associated with blood indices or morphological traits. The correlation of our GWAS results with reported variant associations from flow cytometry, bulk blood traits and high content imaging studies provides orthogonal validation for the ability of BCP to capture meaningful aspects of immune cell biology (28–30).

Compatible with our genetic findings, BCP morphological features were associated with inflammatory endpoints such as asthma, allergies and sinusitis. We also observed morphological associations in cells with elongated morphology to leukocyte counts and healthcare visits related to contraceptive management. However, a more careful examination of this association is warranted as the endpoint does not distinguish between use of specific contraceptives, and includes any contraceptive-related visits to healthcare. Similarly to previously reported age-related decline of mitochondrial activity (7), we found mitochondrial variability in these cells to correlate with donor age.

We anticipate that our extensive dataset will serve as a valuable resource for future research to delineate disease phenotypes by capturing the range of normal morphological variability, and representing a reference for comparative analyses. Our and similar future datasets could establish a baseline for healthy individuals in Cell Painting-based drug screens of pathologies of the blood (46), such as leukemias. Towards this goal, our proof-of-concept application of BCP on AML patient PBMC samples differentiated between leukemia FAB subtypes. The results revealed distinct morphological and cell type-specific differences within AML samples, demonstrating the potential of BCP for profiling leukemic samples. Nevertheless, comprehensive benchmarking with a larger number of samples is essential for establishing the BCP as a reliable tool for AML classification and profiling towards diagnostics.

We foresee the training of label-free models with large BCP datasets as a promising direction for inexpensive morphological profiling (38,47). Our publicly available BCP Atlas offers a foundation for label-free models (48–50) and large-scale biological discovery.

## Methods

### Healthy blood donor sample collection

The healthy blood donor samples profiled in this study originate from a FinnGen project in collaboration with the Finnish Red Cross Blood Service Biobank. The Blood Service Biobank contributed DNA samples from 58,000 blood donors to FinnGen. For a subset of these donors, live-frozen PBMCs, plasma, and serum samples were collected and stored, along with genotype data generated earlier in FinnGen^1^. Full blood count and clinical chemistry analyses were also performed from collected whole blood and serum. All biobank samples from Finnish Red Cross Blood Service Biobank were collected along with normal blood donation from donors who met the blood donation health requirements and who had given biobanking consent (51). For blood donation, blood donors must be between 18 and 70 years of age, and feeling healthy and well. If a medical condition has been diagnosed, it must be well-controlled and medicated. Certain medications and medical conditions may disqualify potential donors from donating blood either temporarily or permanently.

Donated blood is collected in a standard blood bag system with Citrate Phosphate Dextrose (CPD) as anticoagulant (CompoFlow®, Fresenius Kabi, Cat.No. CQ32250). After blood donation, donated blood was transported overnight to the Blood Service’s production facility and processed within 24 hours. The red blood cells and plasma were separated from the donated whole blood by centrifugation and used for preparation of blood products. The remaining Buffy Coat fraction (∼55 ml) was passed for isolation of PBMCs to the Blood Service Biobank. PBMCs were isolated from the Buffy Coat with Ficoll isogradient centrifugation (52,53). After the isolation, PBMCs were stored in a freezing medium (5% filtered heat-inactivated human AB serum (Valley Biomedical), 2 mM L-glutamine (Gibco), and 1% penicillin/streptomycin (10,000 units/mL and 10,000 µg/mL respectively) (Gibco) in RPMI 1640 (Gibco) containing 10% DMSO (WAK-chemie medical GMBH)) in cryotubes (1.9 ml Tri-coded Tube, FluidX 65-7641, Azenta) and transferred into the automated gas phase liquid nitrogen freezer for long-term storing at -180 C until release and transported to the research laboratory.

### Blood Cell Painting protocol ‒ overview of the procedure

Steps involved in the modified Cell Painting protocol for PBMC samples are described below, with a more detailed protocol of the methods found in the **Supplementary Text**.

### Cell culture, staining and fixation of PBMC samples

The media used for thawing and culturing the cells was a 5% filtered heat-inactivated human AB serum (Valley Biomedical), 2 mM L-glutamine (Gibco), and 1% penicillin/streptomycin (10,000 units/mL and 10,000 µg/mL respectively) (Gibco) in RPMI 1640 (Gibco). Thawing of the samples was completed according to standard protocols. The cells were washed twice to remove cytotoxic DMSO and resuspended in culture media to contain approximately 50,000 cells/ml. Prior to the seeding of the cells, the wells of 384-well plates (CellCarrier-384 Ultra/PhenoPlate 384-well, Revvity) were coated with Poly-L-lysine hydrobromide (MP Biomedicals) to enhance electrostatic interactions between the negatively charged ions on the cell membrane to the positively charged coated well (54).

Each imaged plate contained ten donor samples and a control sample. As control samples, we used sister samples from a large cohort of PBMCs collected from donors during a singular blood donation. Samples collected from a singular donor were used on 11–17 plates as controls. With the aim to image at least 100,000 cells per donor, approximately 50,000 cells per well were added in parallel to five wells on 384-well imaging plates. One control sample was added on each plate to five wells, aiming for at least 100,000 imaged cells per control. The cells were incubated at 37°C for 1.5 hours prior to adding Mitotracker (see below). After a complete two-hour incubation time at 37°C and 5% CO_2_, the plate was fixed with 8% paraformaldehyde.

### Fluorescence staining of healthy donor PBMCs

The staining pipeline used was based on an optimized Cell Painting assay (4,5). The mitochondria of the living PBMCs were stained with Mitotracker Deep Red (Invitrogen) before fixation of the cells. Wheat-germ agglutinin/Alexa Fluor 555 (WGA) conjugate stock (Invitrogen), Concanavalin A/Alexa Fluor 488 conjugate stock (Invitrogen), Phalloidin/Alexa Fluor 568 conjugate stock (Invitrogen), SYTO14 green fluorescent nucleic acid stain stock (Invitrogen) and Hoechst 33342 stock (Invitrogen) were added after fixation and permeabilization of the cells, described in detail in **Supplementary Text**.

### High content imaging with Opera Phenix microscope

The plates were imaged with Opera Phenix High Content Screening System (Revvity, Waltham, MA, USA) using 40x water immersion objective (NA 1.1) and four lasers (405 nm, 488 nm, 561 nm and 640 nm). A total of 64 (8x8) fields-of-view (FOV) with 5% overlap were imaged per well using five predetermined Z focus planes (1.5 µm distance between planes) with laser-based autofocusing. The images were captured with two Andor Zyla sCMOS cameras (16-bit, field of view 320x320 μm^2^, effective xy resolution 0.33 µm). Both confocal and widefield images were acquired. Detailed description of excitation and emission sources are described in **Supplementary Table 3**. The experimental workflow was optimized and designed so that a single person could prepare up to four plates during a week (one plate per day), and perform sample staining during the last day of the work week. Plates were then imaged by automated microscopy over the weekend, with one plate taking approximately 10 hours to image. Approximately 35 TB of images were generated during data collection (530 GB/confocal plate, 320 GB/widefield plate).

### Image analysis

Biology Image Analysis Software (BIAS) (https://www.sct.bio/the-bias; Single-Cell Technologies) was used for primary image processing, segmentation, and feature extraction (55). Both confocal and widefield images were processed using the same analysis workflow with the exception that widefield images had one extra channel used in feature extraction (**Supplementary Text**). 3D image stacks were first projected into 2D images using maximum intensity projection for fluorescence images and average intensity projection for brightfield images. Non-uniform illumination was corrected on these images using the CIDRE method (56). Next, nuclear and cellular regions were segmented using two pre-trained deep learning models. Nuclei were segmented using the nucleAIzer (57) model and the cellular regions using the nucleAIzer extension for cytoplasm segmentation (available at: https://zenodo.org/records/6790845). These two segmented regions were linked as cell objects including one segmented region from both models. Classical intensity, shape, and texture features were extracted from all channels on cell object regions to present 186 BCP cellular features.

To ensure that cellular features from different plates are comparable, features were standardized plate-wise with respect to cellular features of the control sample on the plate. First, for each feature, we calculated mean *μ* and standard deviation 𝜎 for control cells. Then, the donor cellular feature values 𝑥 were standardized as 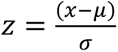.

### Outlier and edge removal

Out of ∼55 million cells imaged, approximately 5 million were from control samples and were excluded from analysis after being used for feature scaling (Methods). Outlier removal was done based on features “NUCLEUS INTENSITY-MEAN Alexa 568” and “NUCLEUS INTENSITY-MEAN Hoechst 33342”, where all cells with scaled features with values over 40 were removed from the dataset. These were mostly falsely segmented high-intensity artefacts. In addition, edge cell removal was done based on values of features “centroid_x”, “centroid_y” and “CELL ENCLOSING CIRCLE RADIUS”, where cells were removed if they were touching the edge of the well to ensure that all analyzed cells fit completely in the field-of-view. These steps resulted in 44.8 million cells for downstream analysis and batch correction.

### Batch correction of BCP cellular features

Three methods were employed for the batch correction of cellular features: Harmony (18), pyComBat (20) and least-squares linear regression. Harmony clusters data points into temporary clusters, computes correction factors based on the cluster centroids, and repeats until satisfactory entropy and diversity in the clusters have been reached. PyComBat is a Python implementation of the ComBat (19) batch correction method, which uses parametric and non-parametric empirical Bayes approaches to correct batch effects. We evaluated three potential sources of batch effect: plate, well, and field-of-view (FOV), and their combinations. For linear regression, corrected features were derived by subtracting the observed feature value from the feature value predicted by the correction model.

We used a batch effect score to evaluate the effectiveness of the above methods by measuring whether cells from the same plate are more clustered together than expected, indicative of batch effect. The score is derived by computing, for each cell 𝑖 in the data, the fraction of cells from the same plate as cell 𝑖 amongst 𝑘 nearest cells, and normalizing by the expected fraction. To find the nearest cells, Euclidean distance in the UMAP space of cellular features was used. For each batch correction experiment, let 𝑁 be the number of cells in the whole dataset, 𝑛^𝑝^ be the number of cells from the plate 𝑝, and the number of cells from the same plate in the nearest 𝑘 neighbors be 𝑥_𝑖_ where 𝑖 is the cell id. We define the batch effect score as

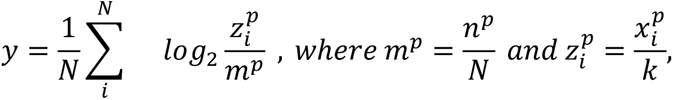

or the average of the logarithm of observed to expected fractions taken over all cells. The batch effect score for an individual cell is zero if the two ratios match, and increases when there are more cells from the same plate than expected, suggesting the presence of a batch effect. The ability of the score to detect batch effects depends on the choice of 𝑘, such that larger values of 𝑘 both allow the detection of smaller batch effects and accommodate larger between-plate differences in cell amounts. The value of k used for this study was 200.

We were not able to perform batch correction on the quality-controlled set of 44.8 million cells due to insufficient memory (1,490 GB RAM) to run Harmony. Therefore, we first ran Harmony on 35 million cells in a single batch, and then corrected the remaining cells as follows: for each remaining cell, feature value was assigned as the average of three Harmony-corrected cells closest with respect to Euclidean distance to the uncorrected cell in the feature space.

### Clustering of cells by BCP cellular features

Clustering of cells by BCP features was done by first performing two-dimensional UMAP (n_neighbors=100, min_dist=0.01) and then applying DBSCAN clustering (58) on the UMAP coordinates. After clustering cells with DBSCAN, all cells not assigned to any cluster were removed as outliers, leaving eight initial clusters. After initial clustering, further subclustering was done based on visual inspection of major clusters that could be clustered further (two largest clusters C1 and C2) (**Fig. 4**). Parameters for subclusters were assessed by testing multiple different values for maximum distance between two points to be considered neighbours (ε) and minimum number of points within ε (minPts). The parameters for initial clustering were ε=0.08 and 35 minimum samples. The parameters for subclustering of cluster C1 were ε=0.1 and 65 minimum samples, and for cluster C2 ε=0.09 and 60 minimum samples.

### Cell typing

To classify single cells into immune subtypes, we used a supervised machine learning pipeline in the BIAS software. We manually annotated ∼200 cells per class to train a Support Vector Machine (SVM) classifier, which was refined using an active learning loop. K-means clustering (iterations = 100, max deviance = 0) was applied to assist in stratified annotation and visualization of the feature space.

Cells were annotated to four categories: monocytes, macrophages, B & T cells, and uncategorised cells. The aim was to maintain class balance during annotation to support robust training and quantification. Annotation criteria were based on established morphologies in multiplexed confocal fluorescence images. Cells were assigned into the four categories as follows (**Fig. 2 A**): 1) monocytes, large cells characterized by a kidney-shaped nucleus (bi-lobed structure). These cells exhibited a bright signal in the ER/Golgi channel (green) and the mitochondrial marker (red). The actin marker (phalloidin) formed a substantial mesh around the nucleus (orange), 2) macrophages, similar to monocytes but with a more rounded nucleus that have differentiated from monocytes in wells, 3) B and T cells, small cells with a rounded nucleus showing low signal intensity in the ER/Golgi (green), mitochondrial (red) and actin channels (orange), and 4) uncategorised, including segmented objects that did not fit the morphologies described above, for instance cell debris, dead cells, and cells exhibiting only nuclear signals. All extracted intensity, texture, and morphology features were used as inputs without prior feature selection, enabling the model to determine optimal feature weighting.

The classifier was evaluated with ten-fold cross-validation: the dataset was partitioned into ten subsets, each in turn used as a test set while training on the remaining data. Classification performance was assessed using a confusion matrix that confirmed high accuracy for B & T cells and moderate separation between monocytes and macrophages. The final classifier model was then trained with the complete training dataset.

### Correlation of cell counts between BCP clusters, and snRNA-seq cell types and blood counts

We compared the proportion of cells in BCP clusters to both snRNA-seq cell type proportions and blood counts with Pearson correlation. For each cell type available in snRNA-seq data (59), we aggregated counts per cell type (“pseudobulk”) and computed the proportion of each cell type for each donor. The snRNA-seq data was released to the FinnGen sandbox in 2023 along with other multiome data (https://docs.finngen.fi/finngen-data-specifics/red-library-data-individual-level-data/omics-data/single-cell-transcriptomics-and-immune-profiling).

Complete blood counts included both L- (percentage) and B-values (count per liter). CBC measures the number and size of red blood cells, white blood cells, and platelets in blood, with hemoglobin and hematocrit levels.

### Flow cytometry analysis

Cell populations were profiled using high-throughput flow cytometry (HTFC; FIMM High Throughput Biomedicine unit, HiLIFE, University of Helsinki) (**Suppl. Table 4**). The cell type markers used were CD45 (leukocyte), CD11c (dendritic cell), CD3 (T cell), CD8 (cytotoxic T cell), CD19 (B cell), CD56 (NK cell) and CD14 (monocyte/macrophage), as well as non-viable cell exclusion dye DRAQ7. Individual cell types for comparison to Cell Painting features were gated following the protocol described in **Supplementary Figure 1**.

### Visualization of BCP cellular features

Cellular features were visualized by reducing feature dimensionality to two with Uniform Manifold Approximation and Projection (UMAP) (60). UMAP is a non-linear dimensionality reduction algorithm that aims to preserve the similarities and dissimilarities between data points while reducing the feature space dimensionality. Input cellular features to UMAP were normalized to zero mean and standard deviation of one. The used UMAP parameters were n_neighbors=100 and min_dist=0.01. In order to plot millions of data points, we used Datashader (Datashader v0.16.1., 2024, April 19, https://datashader.org) library to accurately represent the data distributions by aggregating local data points to avoid overplotting. In UMAP plots, local regions with multiple colors (labels) appear darker than regions with fewer colors. CMplot version 4.5.1 was used to visualize multiple GWAS in the same plot.

### Quantification of cell-cell interactions

To detect and quantify cell-cell interactions, we used the subset of 2,090,581 cells from 20 donors, for which cell typing and clustering data were available. The number of proximal cells, measured as the number of cells present within the interaction radius from the center cell, was computed using the KDTree nearest neighbor algorithm (scikit-learn v.1.1.1). The co-localization of immune cells within a 22-25 µm radius has been associated with specific immune processes and clinical variables, indicating biologically significant interactions between cells (61,62). Therefore, given the average cell enclosing circle radius (36.33 pixels), we set the interaction distance to 146 pixels, equivalent to 22 µm or 2 cell diameters (0.15 µm/pixel image resolution). We also computed a measure of relative enrichment (RE) to determine if certain cell types were more prevalent in the set of proximal cells than expected. Let 𝑅 be the total number of cells within the interaction radius 𝑟, let 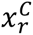 be the number of cells of type 𝐶 within the interaction radius 𝑟, let 𝑁 be the total number of cells in the dataset, and 𝑛^𝐶^ be the total number of cells of type 𝐶. We defined RE as:

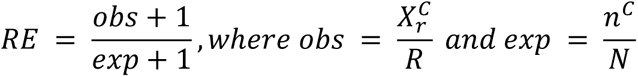

### Principal component analysis of BCP cellular features

Principal component analysis (PCA) was performed for 186 BCP cellular features in order to reduce data complexity and computational burden for subsequent analyses. PCA was done with 20 principal components (PCs) denoted F1–F20 which explained >95% of variance in the dataset. To obtain donor-level features, the average of each PC in each cell cluster was computed for each donor.

### Genome-wide association study for cellular BCP features

Genome-wide association study (GWAS) was performed to find associations between BCP cell features and genotypes in FinnGen (R12). We utilized the FinnGen DF12 dataset with 500,348 donors, 2,502 disease phenotypes and over 21 million genotyped variants (1). We considered 186 BCP features of the four fluorescent channels transformed to 20 PCs (**Suppl. Fig. 10**). GWAS was run for all combinations of the 17 cell clusters and 20 PCs. Cell cluster C7 was not included in the analysis as it contained cells from only 67 donors. The donor-level average value of each PC in each cell cluster was treated as phenotype in GWAS. Altogether 340 GWAS were run (20 phenotypes x 17 clusters). Out of 390 donors imaged, 22 did not have covariate data available or were otherwise excluded from analysis in earlier data collection of FinnGen due to quality control or ancestry (1). The remaining 368 donors had 10.4 million germline variants present with minor allele frequency (MAF) > 0.0125 in FinnGen, and were tested against altogether 22 million germline variants available in FinnGen.

Rank inverse normalization transformation (RINT) was applied to features with inflation (λ>1.05) or deflation of lambda values (λ<0.98). Regenie model (63) was used to run GWAS and the model was trained with a high-quality LD-pruned genetic relationship matrix (GRM) available in FinnGen (1). This GRM contains no population outliers or genetic duplicates, and was further pruned to only contain variants for which BCP donors had MAF>0.0125.

As GWAS covariates, age, sex and the first ten genetic ancestry principal components were used. A base model for associations was also investigated with cellular feature values as independent variable and age, sex and ancestry principal components as covariates.

### Fine-mapping of associated variants in GWAS

Fine-mapping was done for all 30 genomic regions highlighted as significant in GWAS utilizing SuSIE (24) with the FinnGen R12 pipeline and default parameters (max_region_width=6,000,000; p_threshold=5x10⁻⁸; n_causal_variants=10; window_shrink_ratio=0.9; good_cred_r2=0.25).

### Comparing BCP GWAS results to previous studies

We combined summary statistics from previous GWASs with our BCP GWAS by variant rsIDs. For each previous study, the Pearson correlation coefficient *r* was computed between -log_10_(*p*)-values reported in the study and obtained from the BCP GWAS for variant associations (**Suppl. Fig. 11**). Confidence intervals for *r* were estimated using bootstrapping with 1,000 samples.

### Acute myeloid leukemia (AML) patient samples and MOLM-13 cell line sample

To test BCP on PBMCs from leukemia, we used six patient-derived AML (AML1-AML6) samples from five different patients (**Suppl. Table 7**). Two of the AML samples were collected from the same patient, with a ten-month interval between collections. The AML samples were previously clinically subtyped as acute myeloid leukemia with minimal maturation (M1, *n*=3), acute myeloid leukemia with maturation (M2, *n*=1) or acute monocytic leukemia (M5, *n*=2) in the FAB (French-American-British) classification system (64). The two AML M5 samples are from the same patient at diagnosis (AML5) and refractory (AML6). In addition, a MOLM-13 cell line (ACC 554) acquired from the FIMM HTB unit, originally established from the peripheral blood of a patient at relapse of AML (65) (FAB M5a) was used in the study. All AML patient samples and the MOLM-13 sample were thawed, plated, fixed, stained, imaged and analyzed using the same BCP protocol as for the healthy donor PBMC samples. All AML samples were plated together with healthy PBMC samples, aiming at least 100,000 single cells dispersed over five wells, with the standard control sample and imaged with both confocal and widefield microscopy.

### Code availability

The code used to analyze the BCP dataset, and dataset details will be available at https://github.com/bioimage-profiling/blood-cell-painting-paper.

## Supporting information

BCP Additional File 1 - Supplementary Text

BCP Additional File 2 - Supplementary Figures

BCP Additional File 3 - Supplementary Tables

BCP Additional File 4 - Finngen banner

## Data availability

BCP image data generated in this study is publicly available in BioImage Archive (10.6019/S-BIAD2344). Instructions on how to access FinnGen genotype and health registry data are available at https://www.finngen.fi/en/access_results.

## Ethical approval

The Coordinating Ethics Committee of the Hospital District of Helsinki and Uusimaa have approved the FinnGen study protocol Nr HUS/990/2017. A formal biobank decision for the research project has been granted by the Finnish Red Cross Blood Service Biobank. Blood was collected from volunteering blood donors that previously have partaken in a FinnGen study and have signed a biobanking consent form with the Blood Service Biobank (The Finnish Biobank Act 688/2012). The primary cells used in this project were human PBMCs, isolated from venous whole blood, and frozen according to the Blood Service Biobank guidelines and sent to FIMM for thawing, cell seeding, fixation, staining and imaging.

AML samples were collected in a study approved by an ethics committee of the Helsinki University Hospital (Ethics Committee Statement 303/13/03/01/201, latest amendment 7 dated 15 June 2016; latest HUS study permit HUS/395/2018 dated 13 February 2018; Permit numbers 239/13/03/00/2010, 303/13/03/01/2011).

## Author contributions

Blood Cell Painting Assay development: CHS, MP, VP

Blood Cell Painting Assay data modeling: GA, VT, LP, EP

Methodology: CHS, VT, IM, MP, GA, AH, LP, VP, EP

Formal analysis: CHS, VT, GA, CDS, IM

Investigation: CHS, VT, GA, IM, MP, JS, JK, CDS, LU, HMO, LP, VP, EP

Visualization: MP, VT, GA, IM, CDS, CHS

Wetlab: CHS, JH, JS, TR, TA

Data curation: VT, GA, CHS, IM, MP

Software: GA, VT, IM

Resources: JP, AH, JJ, JJM, MA, CAH, RR, MD, AP, LP, VP, EP

Imaging development: AH, MP, CHS, IM

Flow cytometry: TR, TA

Genetic analyses: VT, LU, OK, HMO, EP

Registry data analyses: VT, JK, HMO, EP

Conceptualization, Supervision, Funding acquisition: LP, VP, EP

## Acknowledgements

This study was supported by funding from the Research Council of Finland (322675 and 328890 to EP; 340273 and 346604 to LP), the Research Fund of the Finnish Red Cross Blood Service, FinnGen and iCAN Flagship. The Finnish Red Cross Blood Service Biobank personnel are acknowledged for organizing research project-specific sample collection in blood donation activities. The authors wish to acknowledge CSC – IT Center for Science, Finland, for generous computational resources. HC imaging and analysis were performed at the FIMM High Content Imaging and Analysis HCA unit, and flow cytometry at the FIMM High Throughput Biomedicine HTB unit, both supported by HiLIFE, University of Helsinki, Biocenter Finland, and Euro-Bioimaging. Prof. Peter Horvath is acknowledged for his support in image analysis and for providing BIAS software (Single Cell Technologies Ltd.) for our use. We would like to thank Prof. Matti Pirinen, Pietro Della Briotta Parolo, Vincent Llorens, Dr. Markus Vähä-Koskela, Anna Nylund, and the FinnGen helpdesk for their assistance. Prof. Olli Kallioniemi is acknowledged for scientific discussions. BioRender.com was used in creating figures.

We thank all the donors and patients who have participated in this project by donating their samples.

We want to acknowledge the participants and investigators of the FinnGen study. The FinnGen project is funded by two grants from Business Finland (HUS 4685/31/2016 and UH 4386/31/2016) and the following industry partners: AbbVie Inc., AstraZeneca UK Ltd, Biogen MA Inc., Bristol Myers Squibb Inc. (and Celgene Corporation & Celgene International II Sàrl), Genentech Inc., Merck Sharp & Dohme LCC, Pfizer Inc., GlaxoSmithKline Intellectual Property Development Ltd., Sanofi US Services Inc., Maze Therapeutics Inc., Johnson&Johnson Innovative Medicine Inc., Novartis AG, Boehringer Ingelheim International GmbH and Bayer AG. Following biobanks are acknowledged for delivering biobank samples to FinnGen: Auria Biobank (www.auria.fi/biopankki), THL Biobank (www.thl.fi/biobank), Helsinki Biobank (www.helsinginbiopankki.fi), Biobank Borealis of Northern Finland (https://www.ppshp.fi/Tutkimus-ja-opetus/Biopankki/Pages/Biobank-Borealis-briefly-in-English.aspx), Finnish Clinical Biobank Tampere (www.tays.fi/en-US/Research_and_development/Finnish_Clinical_Biobank_Tampere), Biobank of Eastern Finland (www.ita-suomenbiopankki.fi/en), Central Finland Biobank (www.ksshp.fi/fi-FI/Potilaalle/Biopankki), Finnish Red Cross Blood Service Biobank (www.veripalvelu.fi/verenluovutus/biopankkitoiminta), Terveystalo Biobank (www.terveystalo.com/fi/Yritystietoa/Terveystalo-Biopankki/Biopankki/) and Arctic Biobank (https://www.oulu.fi/en/university/faculties-and-units/faculty-medicine/northern-finland-birth-cohorts-and-arctic-biobank). All Finnish Biobanks are members of BBMRI.fi infrastructure (https://www.bbmri-eric.eu/national-nodes/finland/). Finnish Biobank Cooperative -FINBB (https://finbb.fi/) is the coordinator of BBMRI-ERIC operations in Finland. The Finnish biobank data can be accessed through the Fingenious® services (https://site.fingenious.fi/en/) managed by FINBB.

## Notes

https://github.com/bioimage-profiling/blood-cell-painting-paper

### Competing Interest Statement

The authors have declared no competing interest.

### Summary of Updates

We have added results of an analysis where we correlated cell counts obtained from single-nucleus RNA-seq from the same donors.

