## Supplementary material for "Image-based profiling of healthy donor immune cells with Blood Cell Painting reveals novel genotype-phenotype associations": BCP Additional File 1 - Supplementary Text

#### Protocol for Blood Cell Painting and Phenotypic Analysis

##### Phenotypic imaging of PBMC samples – Blood Cell Painting

###### 5% Human AB serum Cell thawing/Culture media

RPMI 1640 medium, containing 5% filtered (Millex-GP) human AB serum, 2 mM L-glutamine, and 1% penicillin/streptomycin (10,000 units/mL and 10,000 μg/mL respectively).

###### Coating of 384-well plate using Poly-L-lysine

1. Dissolve 50 mg Poly-L-lysine in 5 ml MQ H2O to make 10 mg/ml stock (store in –20 °C).
2. Prepare a working solution containing 1 mg/ml Poly-L-lysine in PBS.
3. Coat the wells with 15 μl working solution.
4. Spin the plate up to 300*g*.
5. Incubate for 5-10 min.
6. Remove excess coating and wash the wells 2 times with 50 μl PBS.
7. Cover the plate with the lid, and store in the refrigerator until plating of cells. Note: Use within 2 weeks

###### Thawing the cells

1. Sterilize your workspace, fume hood and necessary equipment with 70-80% EtOH, before thawing the cells.
2. Prepare enough culture media needed for all samples, approximately 20 ml per PBMC sample.
3. Warm up the thawing media in a 37°C incubator or water bath and prepare a 37°C heat block or water bath for the samples.
4. Mark a 15 ml centrifuge tube for each sample planned to be thawed.
5. Add 10 ml of 37°C thawing/culture media to every centrifuge tube.
6. Retrieve the PBMC samples from liquid nitrogen and place them on dry ice.
7. Thaw the samples in a heat block/water bath, a few samples at the time (~3-5).
8. When the cells start to thaw, spray EtOH over the cryotubes and let the EtOH evaporate in the fume hood.

9. Pipet thawing/culture media carefully back and forth from the cryotube to the centrifuge tube, 2-3 times, and wash out the cryotube with media once at the end.
10. Centrifuge the centrifuge tubes containing the sample for 10 min x 300$g$.
11. Discard the supernatant
    Note: Be careful not to mix the cell pellet.
12. Add 5 ml thawing/culture media to the centrifuge tubes and suspend the pellet with the media.
13. Centrifuge the centrifuge tubes for a second time 10 min x 300$g$.
14. Discard the supernatant and add 1 ml thawing/culture media and suspend it with the cell pallet.
    Note: Be careful not to mix the cell pellet
15. Count the cells using a Bürker chamber or an automated cell counter and make the necessary dilutions.
    Note: Dilute the cells according to your own plate and experiment requirements.
16. Add 25 µl of the diluted cells in cell culture media to a 384-well plate (CellCarrier-384 Ultra/PhenoPlate 384-well, Revvity) to get 50 000 cells per well (2 000 000 cells/ml).
    Note: use a low profile polystyrene lid to reduce evaporation.
17. Leave the cells to recover by incubating the plates with cells for 1.5 hours in 37°C and 5% $CO_2$.

**Live cell staining with Mitotracker**

1. Mitotracker Deep red (M22426): 91µl of DMSO is added to a vial of 50 µg Mitotracker to make a stock solution of 1 mM.
2. Prepare stain solution 1:333 Mitotracker (1 mM) in culture media.
3. After the incubation of PBMCs on the plates (17.) for 1.5h in 37°C, add 5 µl staining solution to each well. Final dilution of mitotracker is 1:2000.
4. Spin the plate in the centrifuge up to 300$g$ to ensure equal distribution of the Mitotracker stain in wells and to remove air bubbles.
5. Incubate the plate for an additional 30 minutes at 37°C and 5% $CO_2$. In total, the incubation time is 2 hours.
6. Continue with PFA fixation immediately after incubation.

**Fixing**

1. Add 30 µl of 8% PFA to the wells (V=30 ul) to get a final 4% of PFA.
   Note: use fume hood.
2. Spin the plate up to 300$g$ to ensure distribution of the added PFA and to remove air bubbles.
3. Incubate at room temperature for 20 min.

4. Remove PFA.
   Note: hazardous waste.
5. Wash by adding 50 µl PBS and centrifuge the plate for 5 min x 300*g*.
6. Remove PBS and add 70 µl PBS to the wells.
7. Store the plate at + 4C°.

**Staining - Cell Painting**

- Hoechst 33342 (H1399) 100 mg: 10 ml sterile H20 is added to create 10 mg/ml stock.
  **Staining solution 2ug/ml (1:5000)**
- Phalloidin /AlexaFluor 568 conjugate stock (A12380): 1.5 ml of methanol is added to one vial of phalloidin containing 300 units.
  **Staining solution 20µl/ml (1:50)**
- Wheat-Germ Agglutinin/ AlexaFluor 555 conjugate stock (W32464): 5 ml of dH2O is added to a vial containing 5 mg product, creating a 1 mg/ml stock.
  **Staining solution 2 ug/ml (1:500)**
- Concanavalin A / AlexaFluor 488 conjugate stock (C11252): 1 ml of dH2O is added to a vial containing 5 mg product, creating a 5 mg/ml stock.
  **Staining solution 100 ug/ml (1:50)**
- SYTO14 green fluorescent nucleic acid stain (S7576): The product contains a 5 mM stock solution.
  **Staining solution 3.33 uM (1:1500)**

**Staining after fixation**

Permeabilization buffer: 0,3% triton-x in PBS

Staining buffer: 1% BSA and 0,1% tween-20 in PBS

Workflow:

1. Prepare the necessary buffers and prepare a staining solution containing Hoechst, Phalloidin, SYTO14, Wheat germ agglutinin and Concanavalin A.
   Note: Complete all steps with no pauses.
2. Aspirate PBS from wells.
3. Add 50 µl permeabilization buffer.
4. Spin down the buffer up to 300*g* to ensure equal distribution of the permeabilization buffer and to remove air bubbles.
5. Incubate for 10 min at room temperature in the dark.
6. Remove the permeabilization buffer and wash twice with 50 µl PBS.
7. Add 20 µl stainmix.
8. Spin down the stainmix up to 300*g* to ensure equal distribution of the stains and to remove air bubbles.
9. Incubate 20 min at room temperature in the dark.

10. Remove stainmix and wash twice with 50 µl PBS.
    Note: Hazardous waste.
11. Add 70 µl PBS
12. Store the plate in +4 C° covered with parafilm.

Note: Reagents used and plate layout are described in **Supplementary Tables 1 and 2**, respectively.

#### High content imaging with Opera Phenix microscope

Imaging is completed using Opera Phenix High Content Screening System (Revvity, Waltham, MA, USA) using 40x water immersion objective (NA 1.1), with four lasers (405 nm, 488 nm, 561 nm, and 640 nm). Plates are imaged with confocal and widefield microscopy. With confocal microscopy, wells are imaged in 5 optical sections: from -2.0 um to 4 um, with 1.5 um distance between the planes. The distance between planes is 1.5 um instead of the recommended 0.5 um to optimize time and cost efficiency. The outer fields of the well are left out and the center 8 x 8 fields are imaged with a 5% overlap. With widefield microscopy, the wells are imaged in 2 planes: 0 um and 2 um, with 2 um distance between planes. The same 8 x 8 fields are imaged with the 5% overlap, but with the advantage that the SYTO14 signal can be separated from the other channels. Details of the channels, stains and excitation and emission sources used in BCP are listed in **Supplementary Table 3**.

#### Image analysis using BIAS software

Biology Image Analysis Software (BIAS) by Single-Cell Technologies (https://www.sct.bio/the-bias) was used for primary image preprocessing and image analysis (Mund, 2022).

1. Illumination correction: All channels are corrected separately using CIDRE method with auto gain and auto zero-light options selected, as well as zero-light preserved option selected.
2. Flatten - fluorescent channels: The module is used to convert 3D fluorescent image stacks into 2D images. MIP (maximum intensity projection) is used as the flattening method to set each output pixel as the highest intensity input pixel in the stack on that position.
   Note: Make sure to select the right input of images.
3. Flatten - brightfield channel: AIP (average intensity projection) is used as the flattening method for brightfield stacks to calculate the mean value of the input intensities for each output pixel.
   Note: AIP is used instead of MIP for the brightfield channel as the content of

brightfield image is represented by contrast instead of fluorophore counts as in fluorescence microscopy.

4. Image filters: The module is used to split the illumination corrected and flattened image into 4 images, with the split rate on the x-axis and y-axis set as 2. The module is used to save GPU memory in the following segmentation steps.
5. Nuclei segmentation: Generic nucleus segmentation (v1.0, the nucleAIzer model) is used as the model to segment nuclei stained with Hoechst 33342, with input spatial scaling set to 1.0 and segmentation window to 2048 px. Detection confidence is set to 20% and contour confidence to 50% with minimum size of 250 px and maximum size of 5000 px.
6. Cell segmentation: Generic cytoplasm segmentation (v1.0) is used as the model to segment the cells from a merged image including Hoechst 33342, Alexa 488 and Alexa 568 stained images, with input spatial scaling set to 1.0 and segmentation window to 2048 px. Detection confidence is set to 1%, and contour confidence to 10% with minimum size of 1000 px and maximum size of 500000 px.
7. Mask operators: The module is used to select cells containing a segmented nucleus within the segmented cell. The masktype is selected as mask2d with the primary input being the output from nuclei segmentation and the secondary input being the output from cell segmentation. The selected objects are relabeled.
8. List creator: The module creates an item list that links segmented nuclei and cell objects together. The inputs are the output from the Nuclei segmentation module and the output from the Mask operators module.
9. Feature extraction: The module is used to calculate shape, texture and intensity features from the cells. The output from the Image filters module is set as the input for image2d and the output from the List creator module is set as the input for items. Nucleus is selected for objects, all channels are included as well as all features.
10. Statistics: The module can be used to acquire SQL queries and visualize the data.
11. Exporters: The Exporters module is used to export BIAS data types as general imaging formats and general tables and/or text formats.

### Flow cytometry

Cell populations were profiled by high-throughput flow cytometry (HTFC).

1. Following thawing of the PBMCs, the cells were suspended to RPMI containing 5% human AB serum, 2 mM L-glutamine, and 1% penicillin/streptomycin (10,000 units/mL and 10,000 µg/mL respectively), and let to recover for 1 hour at 37°C and 5% $CO_2$.

2. PBMCs were then plated to conical bottom 384-well plates (Greiner), 80,000 live cells in 30 µl in each well.
3. Monoclonal antibodies (BD Biosciences, San Jose, CA, USA), CD45 (#560777), CD11c (#565806), CD3 (#563219), CD8 (#347313), CD19 (#345789), CD56 (#560842) and CD14 (#555399), and dead cell exclusion dye DRAQ7 (#564904) were added with Echo 525 (Labcyte Inc., Sunnyvale, CA, USA), and stained for 30 min at room temperature.
4. Cells were analyzed with iQue3 (Sartorius, Germany) HTFC.
5. ForeCyt software (Sartorius, Germany) was used to analyze the remaining viable cells.
6. Briefly, the cell singlets were identified on the basis of FSC-A (forward-scatter area) versus FSC-H, and live cells were identified by excluding DRAQ7-positive cells, followed by identification of leukocytes (CD45+). Further characterization was done for NK-cells (CD56+CD3-), monocytes (CD14), B-cells (CD19), T-cells (SSC-A, CD3 and CD8) and dendritic cells (negativity for other lineages and CD11c+) from the leukocytes.

Work was carried out at FIMM High Throughput Biomedicine unit (HiLIFE, University of Helsinki).

**Gating individual cells for cell types in flow cytometry**

To assign cells with cell types in flow cytometry, we performed the following gating procedure. Marker positivity thresholds were estimated from the intensity distributions for each channel as follows: DRAQ7 = 30000, CD3 = 80000, CD8 = 40000, CD19 = 60000, CD56 = 60000, CD14 = 65000, CD11c = 50000. For monocytes an additional granularity component was added due to their larger size from other cells (SSC, side scatter = 50000). Gating strategy is shown in **Supplementary Figure 1**.
