## Supplementary material for "Image-based profiling of healthy donor immune cells with Blood Cell Painting reveals novel genotype-phenotype associations": BCP Additional File 2 - Supplementary Figures

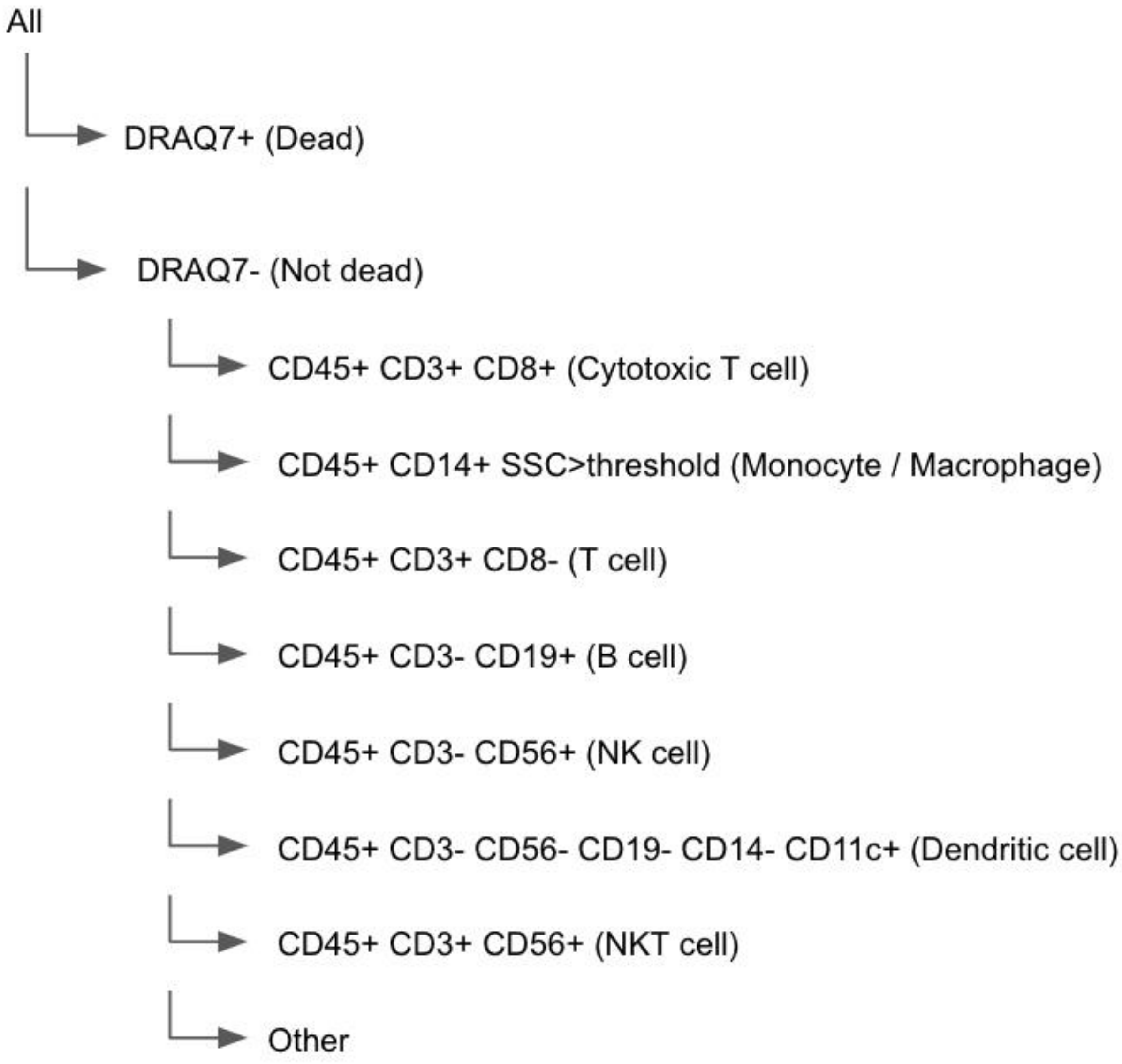


***Supplementary Figure 1***. *Gating strategy of flow cytometry cell typing.*

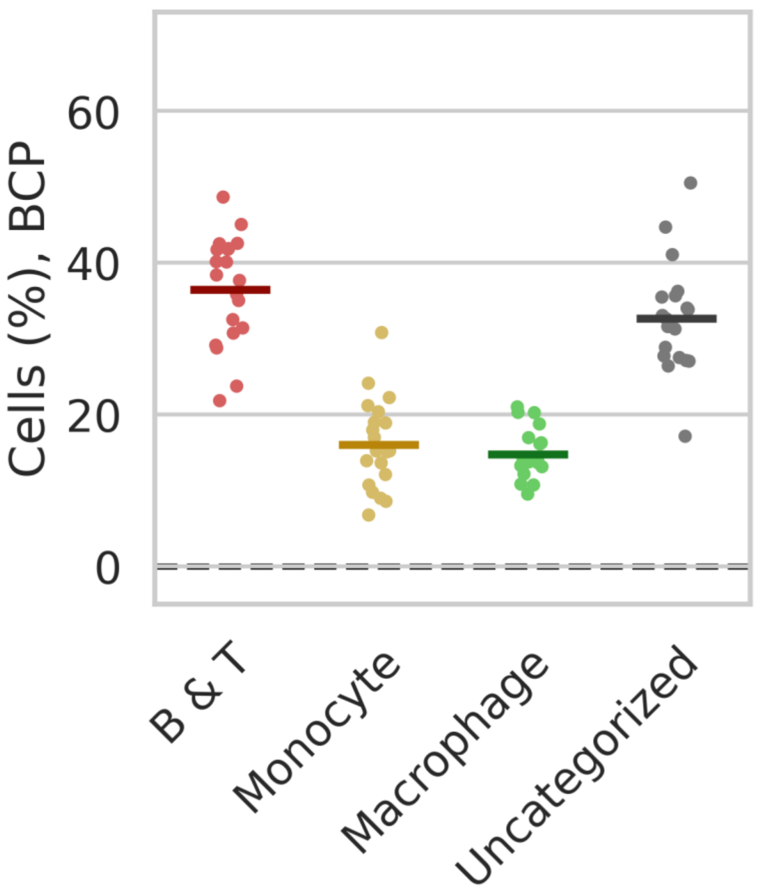


***Supplementary Figure 2***. *Donor-specific proportions of cells by type assigned by the supervised learning procedure in the BCP data based expert annotation. Means indicated with lines.*

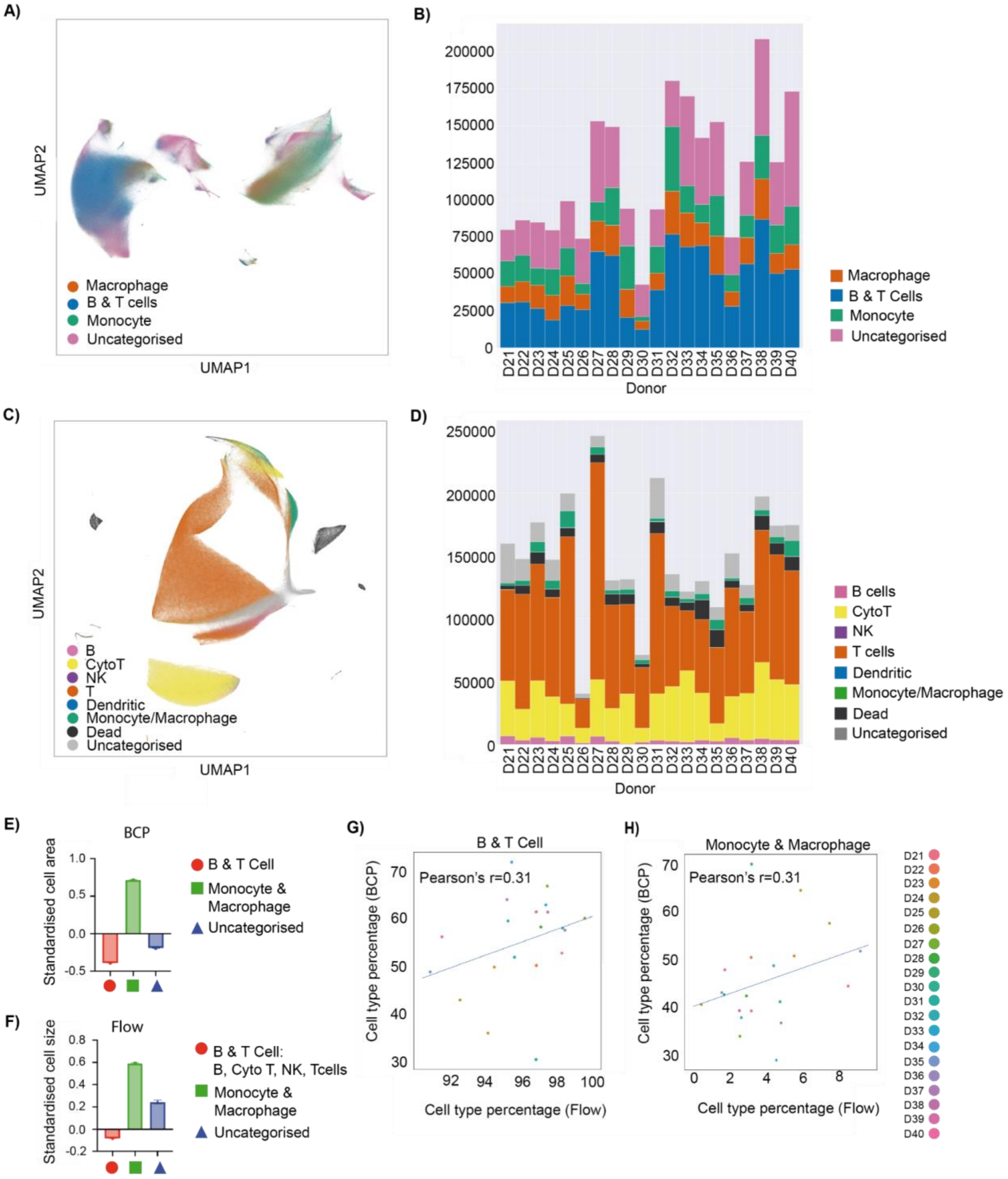


***Supplementary Figure 3**. Comparison of BCP and flow cytometry data for 20 healthy blood donors. A) UMAP representation of the BCP data. B) Frequency of cells by type in the BCP data (cell count on Y-axis). C) UMAP representation of the flow cytometry data. D) Frequency of cells by type in the flow cytometry data (cell count on Y-axis). E-F) Standardised cell areas derived from BCP (E) and flow cytometry (F), respectively. Barplots show a mean with a 95%*

*confidence interval. G) B and T cell, and H) macrophage and monocyte frequencies (%) detected in BCP (Y-axis) and flow cytometry (X-axis), with respect to the total number of B, T, macrophage and monocyte cells.*

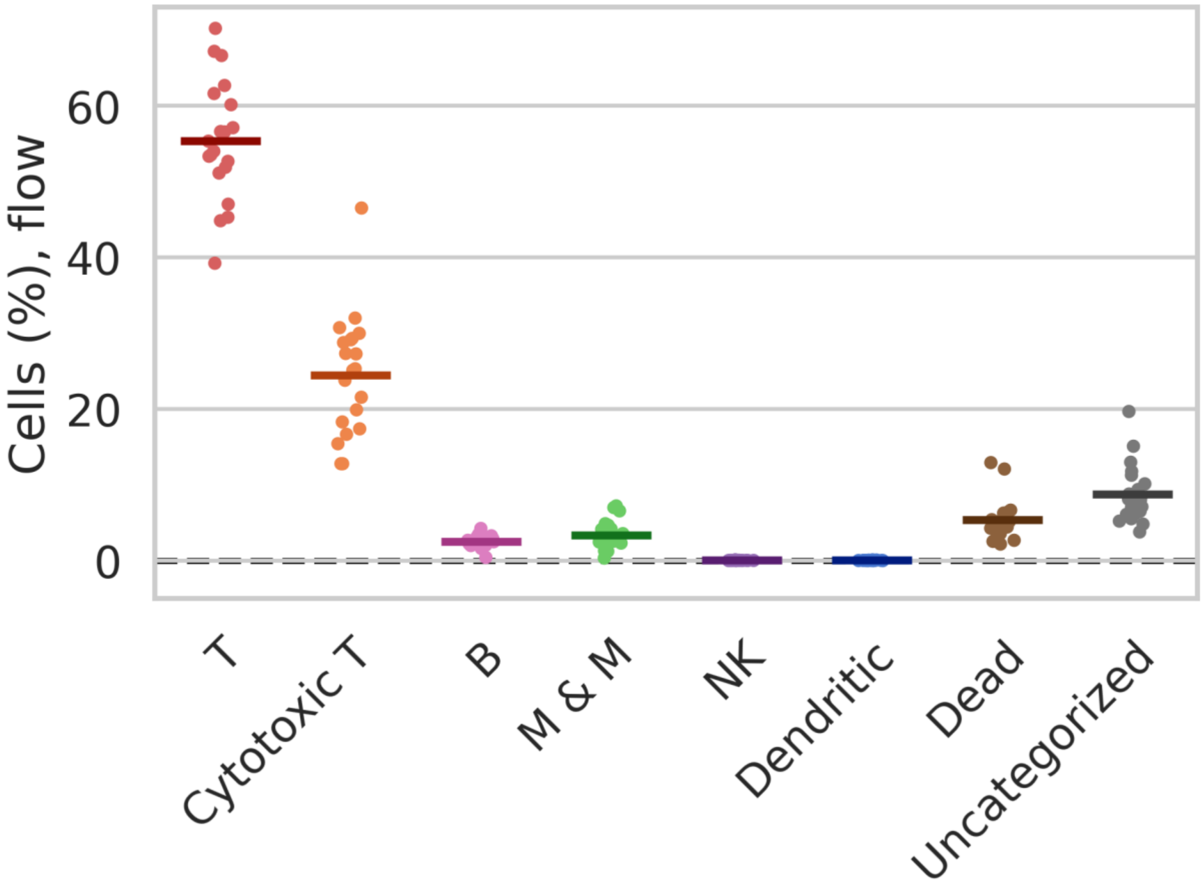


***Supplementary Figure 4****. Donor-specific proportions of cells by type assigned by the gating procedure (Suppl. Fig. 1) in flow cytometry data. Means indicated with lines. M & M, monocytes and macrophages.*

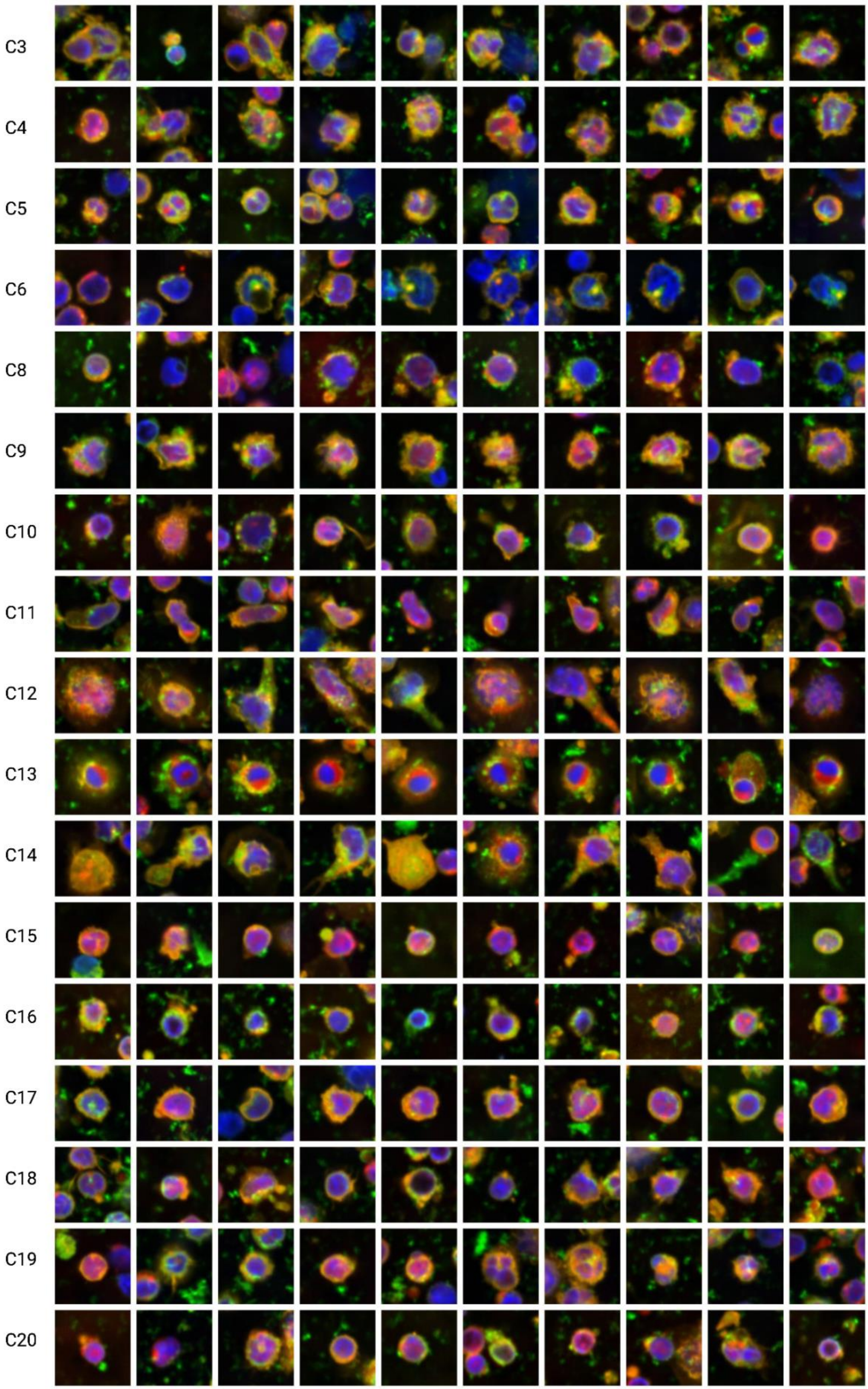
C3
C4
C5
C6
C8
C9
C10
C11
C12
C13
C14
C15
C16
C17
C18
C19
C20

***Supplementary Figure 5.*** *Examples of cell images from each cluster C3-C20 except cluster C7 which was not used in analyses. Created with [BioRender.com](BioRender.com).*

A)

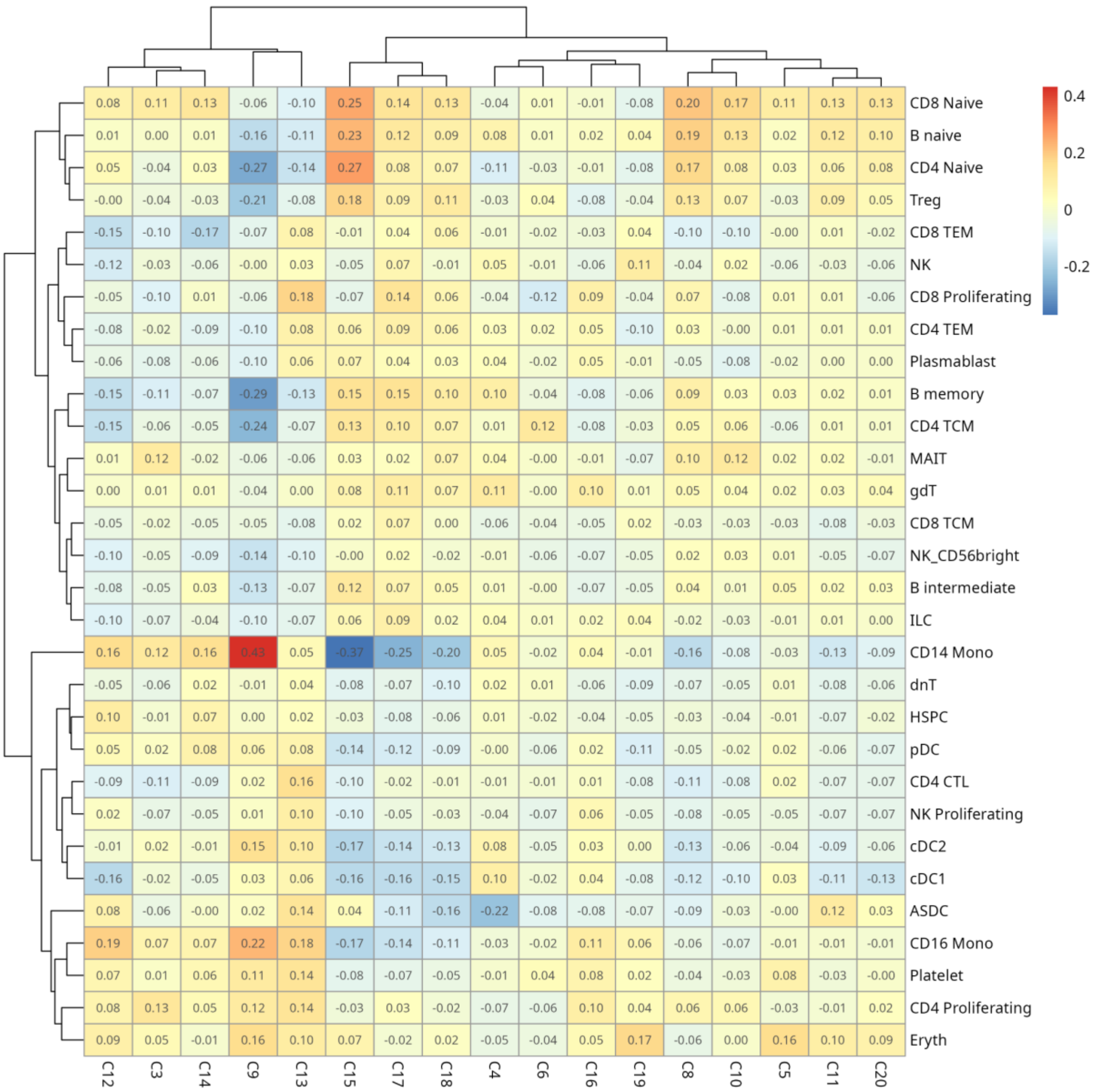


B)

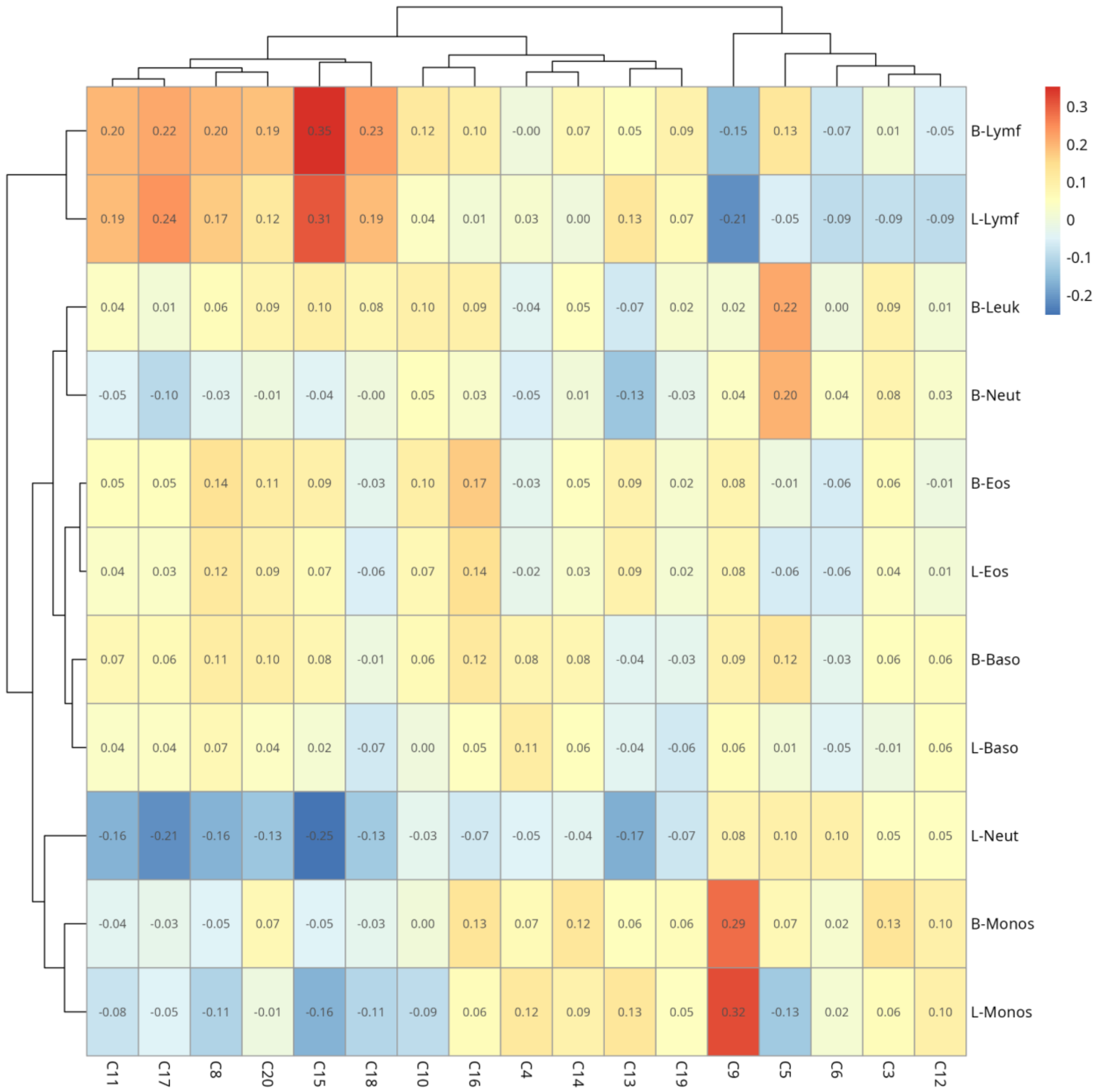


**Supplementary Figure 6.** A) The correlations between snRNA-seq cell type percentages and BCP clusters, B) the correlations between complete blood count values and BCP clusters.

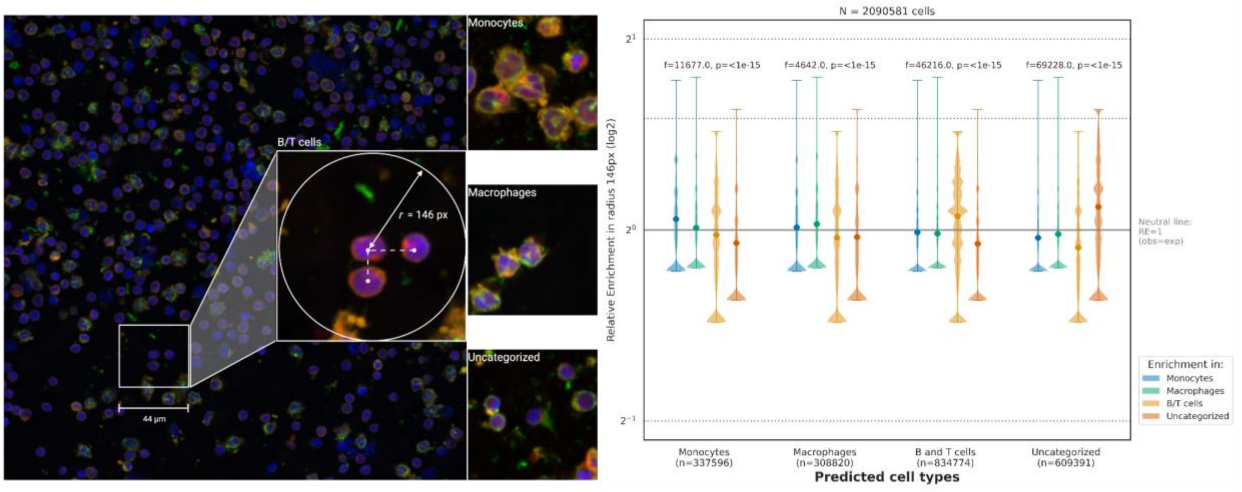


***Supplementary Figure 7.*** *Examples of cell-cell interactions and relative enrichment in monocytes, macrophages, B/T cells, and uncategorized cells. One-way ANOVA F-statistics and p-values suggest significant differences between enrichment in each cell type in monocytes, macrophages, B/T cells, and uncategorized cells. Created with BioRender.com.*

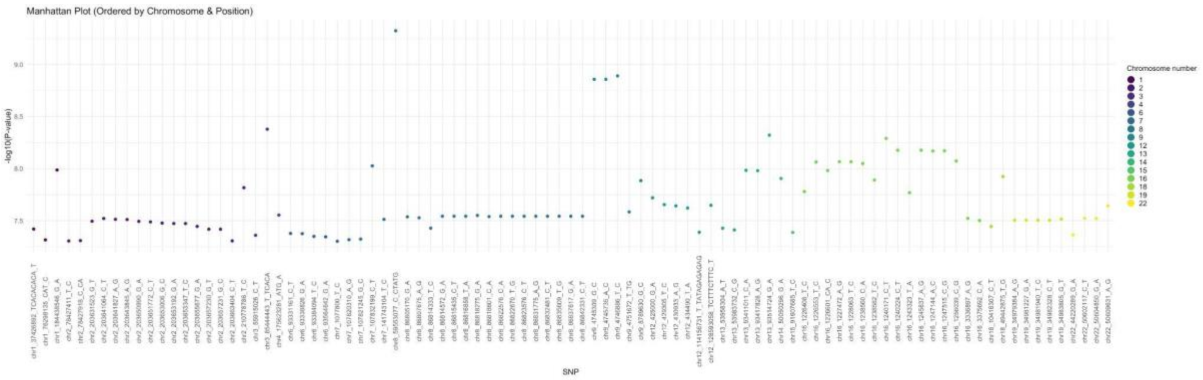


***Supplementary Figure 8.*** *All significant 93 genotype-phenotype associations visualized by their genomic position (X-axis) and -log*$_{10}$*(p)-value (Methods, GWAS QC).*

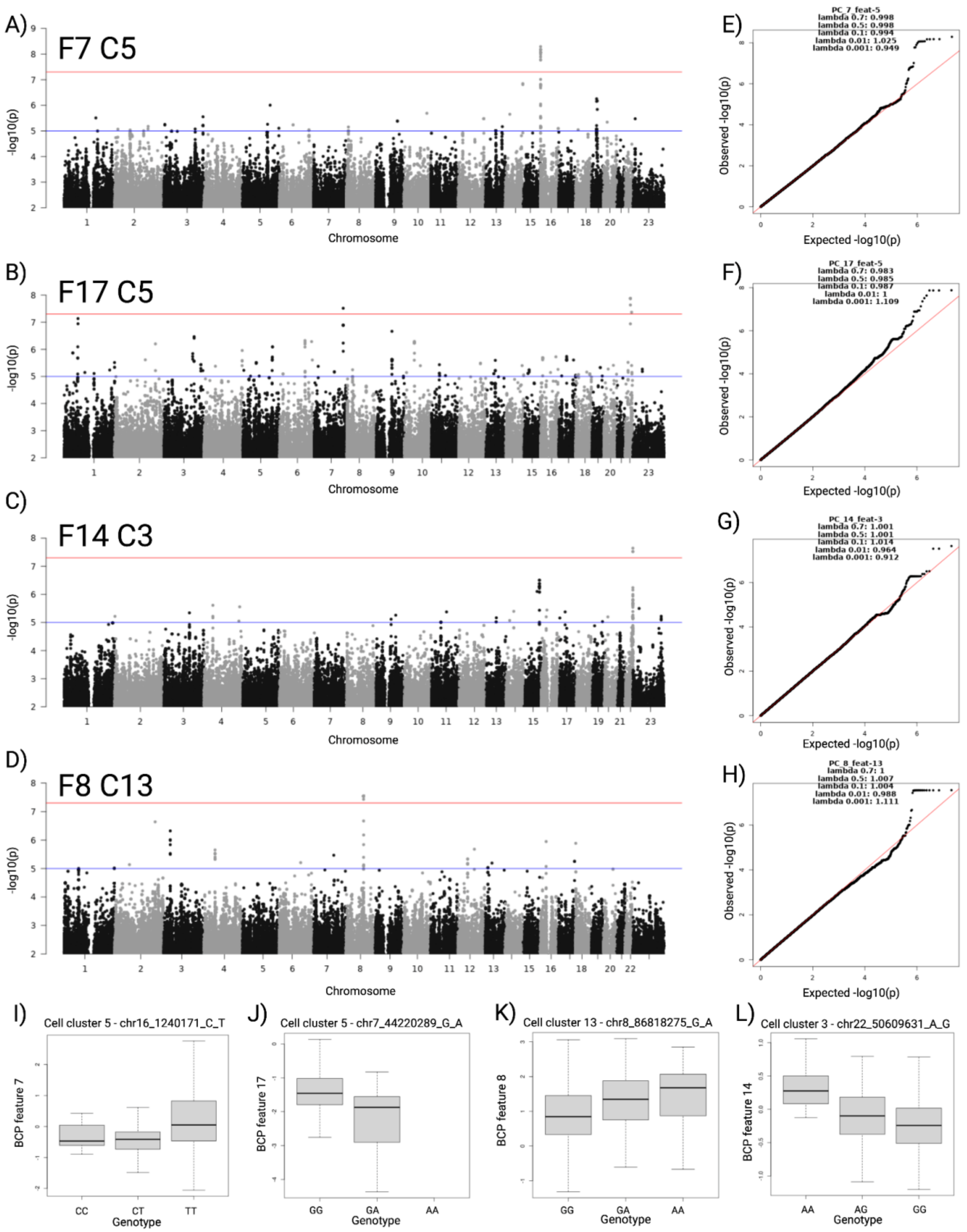


***Supplementary Figure 9**. GWAS results for the top four associating regions between principal components of BCP features and germline variants. A-D) Manhattan plots, E-H) quantile-*

*quantile plots for p-values, and I-L) genotype-phenotype effect plots, where on the x-axis 0 stands for homozygous reference, 1 for heterozygous and 2 for homozygous alternate. Created with BioRender.com.*

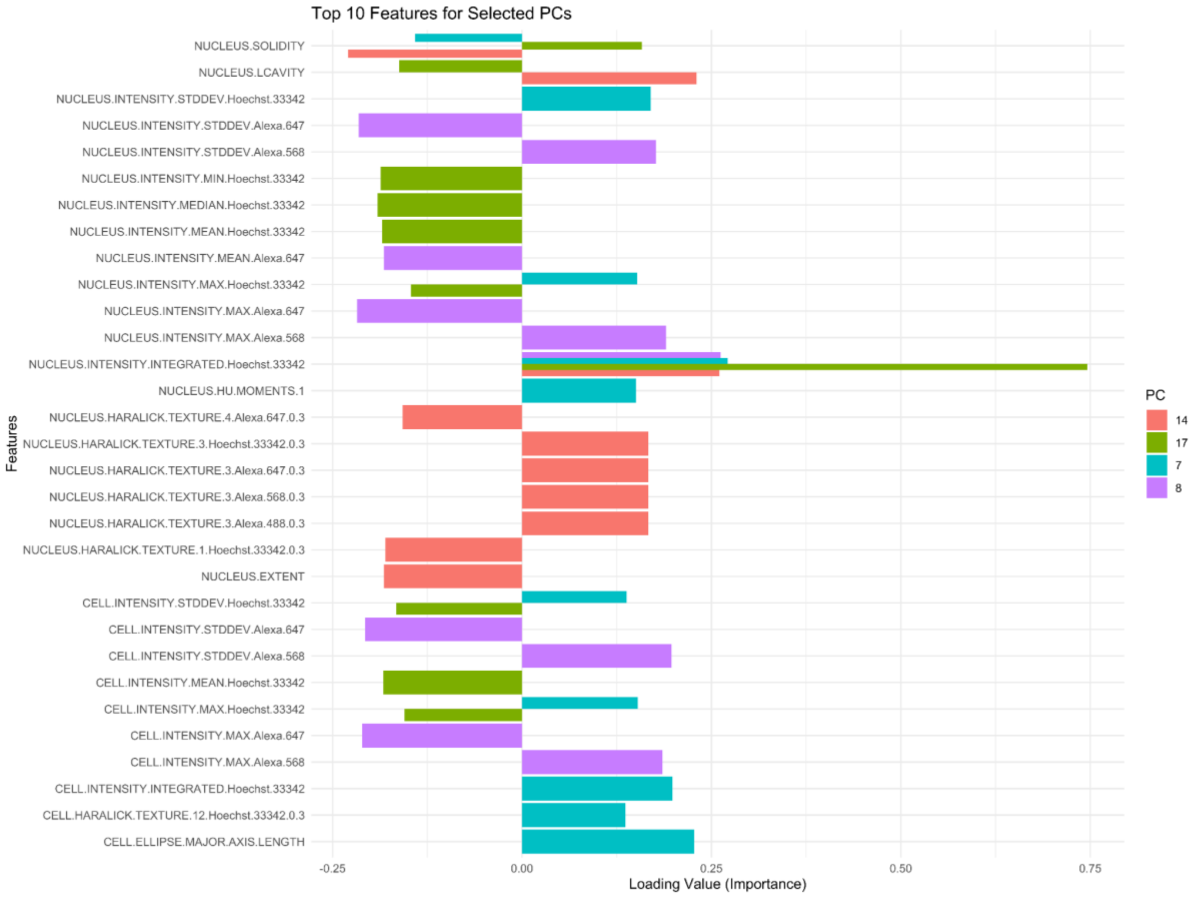


***Supplementary Figure 10**. Principal component loadings for selected four PCs and top ten shared BCP features.*

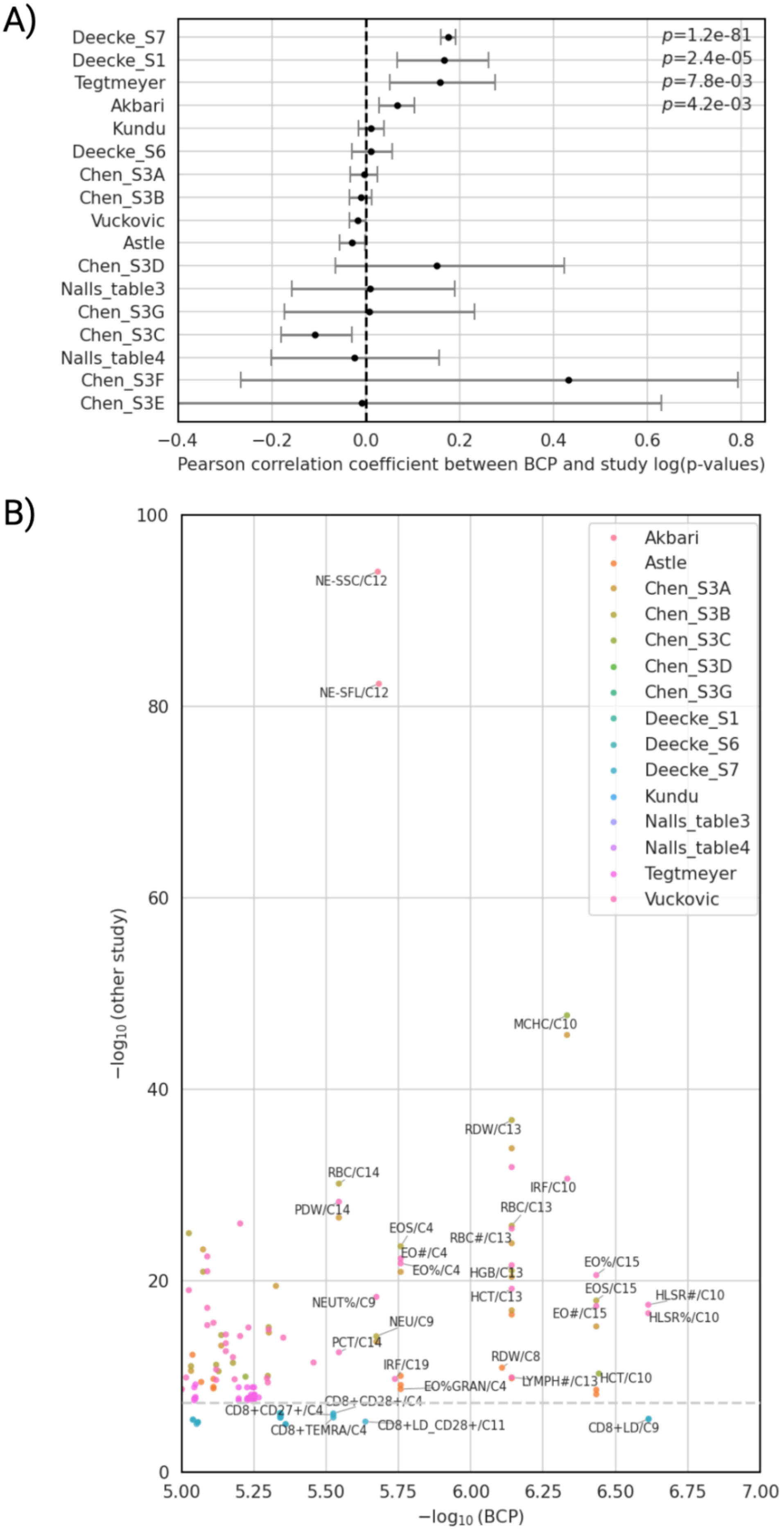
A)
Deecke_S7
Deecke_S1
Tegtmeyer
Akbari
Kundu
Deecke_S6
Chen_S3A
Chen_S3B
Vuckovic
Astle
Chen_S3D
Nalls_table3
Chen_S3G
Chen_S3C
Nalls_table4
Chen_S3F
Chen_S3E
p=1.2e-81
p=2.4e-05
p=7.8e-03
p=4.2e-03
−0.4 −0.2 0.0 0.2 0.4 0.6 0.8
Pearson correlation coefficient between BCP and study log(p-values)
B)
Akbari
Astle
Chen_S3A
Chen_S3B
Chen_S3C
Chen_S3D
Chen_S3G
Deecke_S1
Deecke_S6
Deecke_S7
Kundu
Nalls_table3
Nalls_table4
Tegtmeyer
Vuckovic
NE-SSC/C12
NE-SFL/C12
MCHC/C10
RDW/C13
RBC/C14
IRF/C10
PDW/C14
RBC/C13
EOS/C4
RBC#/C13
EO#/C4
EO%/C4
HGB/C13
EO%/C15
EOS/C15
HLSR#/C10
HCT/C13
NEUT%/C9
EO#/C15
HLSR%/C10
NEU/C9
PCT/C14
RDW/C8
IRF/C19
LYMPH#/C13
HCT/C10
EO%GRAN/C4
CD8+CD28+/C4
CD8+CD27+/C4
CD8+TEMRA/C4
CD8+LD_CD28+/C11
CD8+LD/C9
−log10 (other study)
0 20 40 60 80 100
5.00 5.25 5.50 5.75 6.00 6.25 6.50 6.75 7.00
−log10 (BCP)

***Supplementary Figure 11****. Comparison between variant association results obtained in our study and eight previous studies. A) Pearson correlations and 95% confidence intervals for log(p) values of variant associations reported in both studies. B) a scatterplot with* $-log_{10}(p)$ *values for BCP GWAS (X-axis) and the other GWAS (Y-axis). Labels indicate the trait reported in the other GWAS and the cell cluster (C3-20) reported in the BCP GWAS for selected variants. Colors indicate the other study, and the result table when applicable. Created with BioRender.com.*
